# Mechanical and morphological effects of intervertebral disc injury: a systematic review of in vivo animal studies

**DOI:** 10.64898/2026.03.24.713901

**Authors:** Fangxin Xiao, Jaap H. van Dieën, Andrés Vidal-Itriago, Jia Han, Huub Maas

**Author notes:** Corresponding author: Address: Van der Boechorststraat 7, 1081 BT Amsterdam, The Netherlands.

## Abstract

Intervertebral disc degeneration (IVDD) compromises disc structures and mechanics, yet systematic evaluations of the mechanical responses and their relationship to morphological changes in preclinical models remain limited. This systematic review and meta-analysis synthesized mechanical and morphological alterations following experimental disc injury in in vivo animal models. Searches of MEDLINE, EMBASE and Web of Science databases were conducted in accordance with PRISMA guidelines. Study quality and risk of bias were assessed using modified CAMARADES and SYRCLE tools.

Twenty-eight studies were included. Pooled analyses showed significant reductions in stiffness, Young’s modulus, and disc height, and significant increases in range of motion and degeneration grade, indicating both mechanical and structural deterioration. Young’s modulus appeared to be the most sensitive marker of functional degeneration. By contrast, creep and other viscoelastic responses showed non-significant changes. High heterogeneity was evident across studies, reflecting variability in injury models, species, timepoints, and testing methods. Evidence of publication bias was detected in several domains, and moderate methodological quality was noted with overall insufficient blinding and lack of sample size calculations.

In vivo animal models of IVDD demonstrate robust and consistent mechanical and morphological degeneration after injury. Young’s modulus is a sensitive mechanical indicator, supporting its use in future preclinical research. Standardization of outcome definitions, methodology, and reporting is essential to improve comparability and enhance translation of preclinical findings to clinical research.

## Introduction

Intervertebral disc degeneration (IVDD) is highly prevalent. Population-based MRI imaging studies have reported prevalence of 69.1% in men and 75.8% in women at the L4/5 level^1^, and the presence of IVDD is significantly associated with low-back pain (LBP) ^1^ ^2^. Understanding the pathophysiological mechanisms underlying IVDD is critical for developing effective strategies for prevention, diagnosis, and treatment. IVDD is a complex multifactorial process involving structural, biological, and mechanical alterations^3^ ^4^. A vicious circle may exist among these interdependent factors. For instance, altered mechanical loading of the IVD may alter disc cell homeostasis and extracellular matrix turnover, which may accelerate structural degeneration^4^ ^5^. In turn, degeneration induced changes in disc composition and geometry may further modify load distribution and mechanical behavior, reinforcing the degenerative cascade^4^ ^6^.

Although human studies offer important epidemiological and clinical insights, they are inherently limited in their ability to establish causality. Direct in vivo investigation of the human IVD is further restricted by ethical and regulatory restrictions. Consequently, numerous animal models have been developed in attempts to achieve a comprehensive understanding of IVDD pathogenesis. A wide range of in vitro, ex vivo, and in vivo animal models have been developed^7–9^. Although in vitro and ex vivo IVDD models provide greater control of experimental conditions and could provide insights into isolated aspects of the degeneration processes, in vivo models likely mimic the complex biomechanical and physiological environment of the human spine more closely.

Previous studies have provided validation for the use of quadrupeds as in vivo model for spine research. The normalized geometric parameters (disc height, width, and NP area) of rodent lumbar discs are the closest geometrical representation of the human lumbar discs^10^. And the geometry normalized compression and torsion mechanical properties of the IVD in rat, mouse, rabbit, calf, baboon, and goat represent the mechanical properties of the human lumbar spine, as do the torsion mechanics of the tail discs of rat, mouse, and cow^11–13^.

Current widely used in vivo IVDD animal models can be broadly defined under four categories^9^ ^14^ ^15^: (1) mechanical: (a) physical disruption to the IVD (e.g., stab lesion, needle puncture, nucleotomy), (b) mechanical loading (e.g., static, cyclic compression); (2) biochemical: injection of digestive enzymes (e.g., Chondroitinase ABC, chymopapain) or inflammatory cytokines (e.g., TNF-alpha) that induce destruction of the extracellular matrix (ECM); (3) combination of mechanical and biochemical methods (e.g., mechanical loading or physical disruption with extra biochemical substance injection to the IVD); and (4) spontaneous: (a) aging, (b) genetic mutation or knockout. Among the above methods, structural impairment models based on direct injury to the IVD, including physical disruption and chemical injury (AF or endplate with or without extension to NP), are most commonly used, accounting for 82% of all models^9^. These models offer high reliability, reproducibility, cost-effectiveness, precise control over injury type, location, and development speed^16^. Among these models, needle puncture is the most used (25%) approach used for mechanically induced IVDD, degradative enzymes (e.g., chymopapain, chondroitinase ABC) are the most common used substances for chemically induced IVDD^9^ ^17^.

Most preclinical in vivo IVDD animal models have focused on the gross morphology, histology, and immunohistochemistry following IVD injury, using diagnostic imaging (e.g., radiography (X-ray), MRI) to assess structural alterations, staining techniques (e.g., haematoxylin and eosin, alcian blue/picrosirius red) to reveal histological changes and quantitively assess degeneration degree, biochemical assessments to identify metabolites or changes in protein expression^9^ ^15^. However, only approximately 10% of the included studies have investigated the mechanical responses of the IVD following injury^9^ ^18^. Structural deficits of the IVD (e.g., loss of nuclear material and water content) could alter its mechanical properties, which may lead to changes in load bearing which may in turn affect IVD cell metabolism^6^ ^19^ ^20^. Therefore, biomechanical measurements are essential to improve the understanding of IVD disorders and repair.

Some in vivo IVDD animal model studies demonstrated that IVD injury can result in reductions in stiffness and increases in range of motion (RoM) of the spinal segments^5^ ^15^ ^21–23^, while others showed no significant mechanical alterations in relation to IVD structural deficits^24^ ^25^. The literature reveals inconsistency in reported mechanical results, and there is a lack of systematic comparisons of IVD mechanical alterations across IVDD models, species, and damaged level and structure. Therefore, the primary aim of this study was to establish magnitude and direction of the effects of IVD injury on its mechanical properties. The secondary aim was to explore the relationship between changes in morphological and mechanical properties following IVD injury.

## Methods

This systematic review was conducted in accordance with a protocol registered in the International Prospective Register of Systematic Reviews (PROSPERO), registration number CRD42025630129. The systematic review adhered to the Preferred Reporting Items for Systematic Reviews and Meta-Analyses (PRISMA) guidelines^26^. Ethical approval was not required for this study.

### Search Strategy

A comprehensive systematic literature search was developed in collaboration with a medical information specialist. The databases Ovid/MEDLINE, Ovid/Embase, and Clarivate/Web of Science (Core collection) were searched from inception up to March 14, 2025. The search strategy combined controlled vocabulary and free-text terms, including synonyms and closely related terms for “intervertebral disc”, “spine”, “biomechanical properties”, “injury”, and “animal models”. No restrictions were applied regarding language or publication date. The full search strategies used for each database are detailed in supplementary S1. Duplicate articles were excluded by the medical information specialist using Endnote X20.6 (Clarivate™), following the Amsterdam Efficient Deduplication - AED method^27^ ^28^.

### Inclusion and Exclusion Criteria

#### Inclusion

We only considered studies that reported IVD mechanics in in vivo animal IVD injury models. Models with structural disruption of the IVD were included, including both physical disruption of the IVD (e.g., stab lesion, needle puncture, nucleotomy) and chemical induction of damage (e.g., digestive enzyme injection). Only animal studies written in the English language were included.

#### Exclusion

Studies in which IVD injury was induced post-mortem or the animal had other diseases apart from IVD injury or degeneration were excluded.

### Article Selection and Data Extraction

Article screening was performed by two independent reviewers. Reviewer one screened all the identified reports based on titles and abstracts, followed by full-text screening of eligible articles for final inclusion. Reviewer two validated article screening by 10% of all the identified reports. Data extraction from the eligible studies was performed by two independent reviewers. Reviewer one performed data extraction for all eligible studies, reviewer two performed data extraction validation for 10% of the eligible studies. Disagreements between the reviewers were resolved after discussion, and consensus was eventually reached on all extracted information.

The following data were extracted from all included articles: first author’s name, publication year, animal species, sex, intervertebral disc (IVD) injury model, spine level, interventions in the control group, post-injury timepoints, outcomes of interest (including IVD mechanics and morphological parameters such as disc height, disc height index, and degeneration grade), means and standard deviations or medians and interquartile range (IQR), and sample size. When a control group served more than one experimental group, the total number of control animals was corrected in the meta-analysis by dividing by the number of experimental groups served.

### Risk of Bias and Quality Assessments

The modified Collaborative Approach to Meta-Analysis and Review of Animal Data from Experimental Studies (CAMARADES) checklist^29^ and the Systematic Review Centre for Laboratory Animal Experimentation (SYRCLE) risk-of-bias (RoB) tool^30^ were used to assess the quality of the eligible studies. The SYRCLE RoB results was visualized by using the risk-of-bias visualization tool^31^. A 10-point item CAMARADES checklist (each item received 1 point) was used to evaluate the quality of each individual study:(1) peer-reviewed publication, (2) blinding, (3) sample size, (4) control group(s), (5) animal welfare regulations, (6) animal model, (7) timepoints of outcome assessment, (8) primary and secondary outcomes, (9) statistical analysis, and (10) conflict of interest.

Reviewer one assessed the quality of evidence for all included studies and reviewer two assessed 10% of the eligible studies to validate the assessment of the first reviewer. Discrepancies were resolved by discussion or consensus with a third reviewer.

### Data Synthesis and meta-analysis

Quantitative syntheses were conducted when sufficient data were available (at least 3 studies per domain). To ensure methodological consistency and interpretation, outcomes were grouped into six mechanical domains (stiffness, Young’s modulus), range of motion (RoM), viscoelasticity, creep-strain, and pressure-leakage) and two morphological domains (disc height, degeneration grade). Each domain included a set of relevant biomechanical parameters, for instance, stiffness included neutral zone (NZ) stiffness and directional stiffness measures. A full list of included parameters per domain is provided in Table 1.

**Table 1.** Definition of outcome domains and included parameters.

| Domain | Included Parameters |  |
| --- | --- | --- |
| Mechanics |  |  |
| stiffness | NZ stiffness, stiffness*, torque range (Nm), 3-point bending force |  |
| Young’s modulus | NZ Young’s modulus, Young’s modulus*, pressure Young’s modulus, torque range (MPa) |  |
| RoM | NZ RoM, RoM*, displacement |  |
| viscoelasticity | expected increase after injury: | strain, creep rate, creep displacement, residual deformation |
|  | expected decrease after injury: | creep-derived stiffness (elastic, fast, slow responses; early, late damping stiffness), dynamic moduli (loss, storage, viscous modulus), time constant (fast, slow, early, late, relaxation), damping constant (elastic, viscous), hysteresis* |
| pressure-leakage | pressure (leakage, saturation), volume (leakage, saturation) |  |
| Morphology |  |  |
| disc height | disc height, disc height index |  |
| degeneration grade | histology or MRI based score (e.g., H&E, Safranin O, AB&PR staining) |  |

*NZ, neutral zone; RoM, range of motion; \*includes measurements in various loading directions, such as compression-tension, flexion-extension, lateral bending, rotation; H&E, hematoxylin and eosin staining; AB&PR, Alcian Blue and Picrosirius Red staining*

To account for possible heterogeneity across studies, random-effects meta-analyses were conducted using the standardized mean difference (SMD) and corresponding 95% confidence intervals (CI) to estimate the effect of IVD injury. To harmonize effect directions across parameters in the viscoelasticity domain, effect sizes for outcomes expected to increase following IVD injury were multiplied by −1 after SMD calculation, ensuring a consistent interpretation of effect directions across parameters. Between-study heterogeneity was assessed using the I^2^ statistic, with I^2^ larger than 75% interpreted as substantial heterogeneity^32^. Prediction intervals were also reported to reflect the expected range of effects in future studies. Spearman correlation coefficients (ρ) was calculated to explore the relationship between SMDs of morphological and mechanical properties. Studies employing designs with adjacent IVDs as controls were excluded from the correlation analysis, because anatomical mismatches between measurement levels^33^ could bias the assessment.

To explore sources of heterogeneity, moderator analyses were performed. Univariable meta-regressions investigated both categorical and continuous variables, including species, sex, injury model, spine level, timepoints of mechanical testing following IVD injury, damaged structure, outcome measure, testing direction, and publication year. Moderators identified as significant in univariate analyses were further examined in multivariate meta-regression models. Where appropriate, subgroup analyses were performed based on moderator levels to further investigate variation and estimate pooled effects within specific categories (e.g., injury model, damaged structure). Moderator categories represented by only a single study or level were excluded from multivariate and subgroup analyses to avoid instability and limited generalizability.

All statistical analysis were performed in RStudio (v2025.05.1), with the significant level set at 0.05. Publication bias was assessed using Egger’s test. Sensitivity analysis was performed to examine the robustness of the pooled results. The *influence()* and *leave1out()* functions were used to identify influential studies based on standardized residuals, Cook’s distance, and hat values. Identified influential studies were excluded, and the meta-analysis was repeated to assess the robustness of the effect estimates.

## Results

### Study Characteristics

A total of 4678 studies were found, of which 50 were included for full-text assessment after initial screening. Data from 28 studies were extracted for meta-analysis (Figure 1). Detailed sample characteristics, interventions used to create IVDD models, and main outcome measures of the included studies are shown in Tables 2-3.

**Figure 1.**
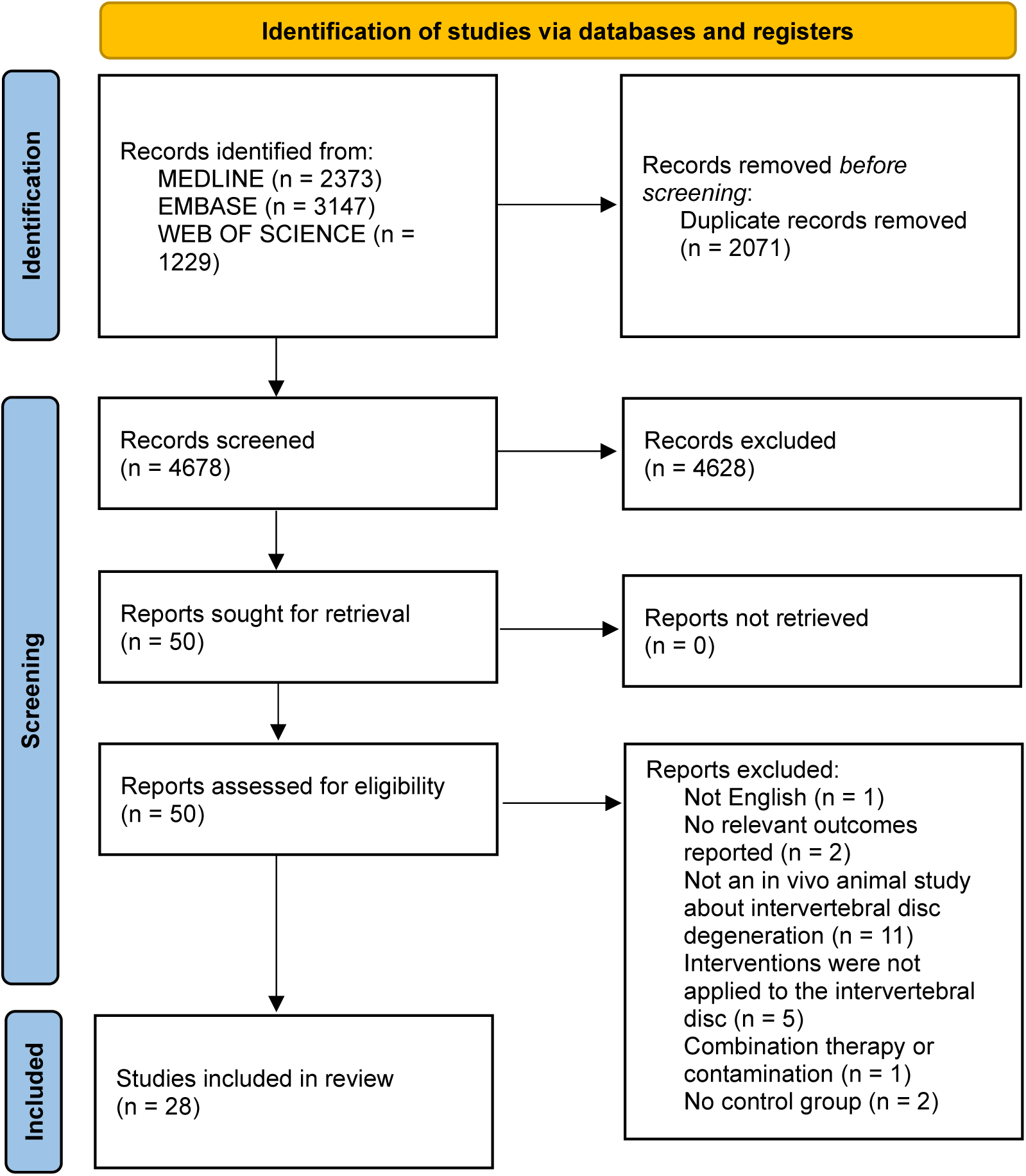
PRISMA diagram of the literature search and eligibility screening

**Table 2.** Characteristics of Included Studies Reporting Mechanical Outcomes.

| ID | Author (Year) | Species (Sex) | Age (Body Weight) | Spine Level | IVDD model | Intervention | Injection Volume / Injury Size | Damaged Structure | Timepoints (weeks) | Reported Outcomes | Groups (n) | Test Direction |
| --- | --- | --- | --- | --- | --- | --- | --- | --- | --- | --- | --- | --- |
| 1 | Boxberger (2008)<sup>55</sup> | rat (male) | 7-9 month (523±43 gram) | lumbar | biochemical | CABC | 1µL (0.25 U/mL), 2.5µL (0.1U/mL) | NP | 4, 12 | NZ modulus, tensile modulus, RoM, creep strain | control (intact:13, sham: 13); injury1 (13); injury 2 (13) | compression-tension; compressive creep |
| 2 | Buser (2011)<sup>45</sup> | porcine (unknown) | 4.6±1.6 year (unknown) | lumbar | physical disruption | nucleotomy | 0.75ml NP tissues was removed | NP | 3, 6, 12 | pressure modulus, leakage pressure | control (5); injury (5) | intradiscal |
| 3 | Calvo-Echenique (2018)<sup>51</sup> | rabbit (unknown) | unknown (5.5-6.5 kg) | lumbar | physical disruption | needle puncture | 18G, 35-40mm depth | AF+NP | 12, 24 | storage moduli, loss moduli | control (4); injury (8) | compression-tension |
| 4 | Colloca (2012)<sup>40</sup> | ovine (unknown) | 24 month (control: 46.9±4.1 kg; injury: 46±4.3 kg) | lumbar | physical disruption | stab lesion | nr.15 blade, 5mm depth | AF | 24 | stiffness | control (6); injury (6) | posterior-anterior |
| 5 | Crowley (2024)<sup>49</sup> | rabbit (female) | 6 month (unknown) | lumbar | physical disruption | fenestration | 1.5mm × 2 mm | AF+NP | 6, 12 | RoM, NZ RoM | control (6); injury 1 (4); injury 2 (6) | axial rotation; flexion-extension; lateral bending |
| 6 | Fainor (2023)<sup>50</sup> | rabbit (male) | 3 month (3 kg) | lumbar | physical disruption | needle puncture | 16G, 21G; 5mm depth | AF | 4, 10 | modulus, RoM, creep displacement | control (3); injury 1 (3); injury 2 (5); injury 3 (5) | compression-tension; compressive creep |
| 7 | Fazzalari (2001)<sup>41</sup> | ovine (male) | 2 year (56±6 kg) | lumbar | physical disruption | needle puncture, concentric tear | 27G | AF | 0, 4, 12, 24, 48, 72 | stiffness | control (4); injury (48) | flexion; left bending |
| 8 | Gullbrand (2017)<sup>37</sup> | goat (male) | ~3 year (unknown) | lumbar | physical disruption, biochemical | nucleotomy; CABC | 22G; 200µl | NP | 12 | modulus, RoM, early / late damping stiffness, strain | control (5); injury 1 (5); injury 2 (5) | compression-tension; compressive creep |
| 9 | He (2022)<sup>53</sup> | rat (male) | 3 month (400±40 gram) | tail | physical disruption | needle puncture | 21G, depth: fully penetrated the IVD | AF+NP | 2 | elastic modulus | control (10); injury (10) | intradiscal |
| 10 | Kaigle (1997)<sup>47</sup> | porcine (unknown) | 4 month (50-60 kg) | lumbar | physical disruption | stab lesion | width 1.2cm | AF; AF+NP | 12 | RoM | control (10); injury 1 (7); injury 2 (7) | flexion-extension |
| 11 | Kaigle (1998)<sup>46</sup> | porcine (unknown) | 6-8 month (80-95 kg) | lumbar | physical disruption | stab lesion | no.11 blade, 15mm depth | AF | 12 | stiffness | control (6); injury (6) | compression |
| 12 | Keller (1990)<sup>48</sup> | porcine (unknown) | 4-8 month (65-80 kg) | lumbar | physical disruption | stab lesion | injury 1: no.11 blade, depth 15mm; injury 2 (partial AF removal): 3.6mm diameter, 15mm depth | AF | 0 | modulus, elastic / viscous damping constant, relaxation time constant, creep rate | control 1 (5); control 2 (5); injury 1 (5); injury (5) | compression-tension; compressive creep |
| 13 | Latham (1994)<sup>42</sup> | ovine (male) | 2 year (unknown) | lumbar | physical disruption | stab lesion | 10mm width, 4mm depth | AF | 0, 24 | stiffness | control 1 (6); control 2 (6); injury 1 (8); injury (6) | axial rotation; flexion-extension |
| 1<br>4 | Leckie<br>(2012a)<sup>44</sup> | porcine<br>(male) | unknown (35<br>kg) | lumbar | biochemical | CABC | 200μL (0.5 U/mL) | NP | 4 | stiffness, NZ<br>length, RoM | control (6);<br>injury (4) | compression,<br>axial rotation,<br>flexion-extension,<br>lateral<br>bending |
| 1<br>5 | Leckie<br>(2012b)<sup>52</sup> | rabbit<br>(female) | 1 year (5 kg) | lumbar | physical<br>disruption | needle<br>puncture | 16G, 5mm depth | AF+NP | 12 | stiffness | control (6);<br>injury (7) | compression,<br>compressive<br>creep |
| 1<br>6 | Lin (2015)<sup>43</sup> | porcine<br>(unknown) | 8 month<br>(32±4.8 kg) | lumbar | physical<br>disruption | needle<br>puncture | 16G, depth: till<br>middle of NP | AF+NP | 0, 1 | leakage /<br>saturation<br>pressure,<br>leakage /<br>saturation<br>volume | control (8);<br>injury 1<br>(8);<br>injury 2 (8) | intradiscal |
| 1<br>7 | Lü (1997)<sup>34</sup> | dog (male) | 1 year (10.3-13.1 kg) | lumbar | biochemical | CABC,<br>chymopapain | chondroitinase<br>ABC: 2.5U/20μL,<br>5U/20μL;<br>chymopapain:<br>120pKU/20μL | AF+NP | 1 | NZ, RoM | control (8);<br>injury 1<br>(16);<br>injury 2<br>(16);<br>injury 3 (8) | axial rotation;<br>flexion-extension;<br>lateral<br>bending |
| 1<br>8 | Martin<br>(2013)<sup>21</sup> | mouse<br>(unknown) | 7.5-9 month<br>(32.7±5.1<br>gram) | tail | physical<br>disruption | needle<br>puncture | 29G, depth: went<br>through NP | AF+NP | 8 | stiffness, RoM,<br>torque range,<br>early / late time<br>constant, creep<br>displacement | control (5);<br>injury 1<br>(5);<br>injury 2 (5) | compression,<br>compressive<br>creep, axial<br>rotation |
| 1<br>9 | Melrose<br>(2012)<sup>23</sup> | ovine<br>(male) | 3-4 year<br>(unknown) | lumbar | physical<br>disruption | stab lesion | 20mm width,<br>6mm depth | AF | 12 | RoM, NZ | control (6);<br>injury (7) | axial rotation;<br>flexion-extension;<br>lateral<br>bending |
| 20 | Mosley (2020)<sup>25</sup> | rat (male, female) | 4 month (unknown) | lumbar | physical disruption + biochemical | needle puncture + TNFα injection | 26G, 3mm depth, triple puncture; TNFα: 2.5 μL (0.1 ng/μL) | AF+NP | 6 | stiffness, RoM, hysteresis, creep displacement, fast / slow time constant, torque range | control (12); injury (11) | compression-tension, compressive creep, axial rotation |
| 21 | Mosley (2019)<sup>54</sup> | rat (male, female) | 4 month (unknown) | lumbar | physical disruption + biochemical | needle puncture + TNFα injection | 26G, 3mm depth, single or triple puncture; TNFα: 2.5 μL (0.1 ng/μL) | AF+NP | 6 | stiffness, RoM, hysteresis, NZ length, creep displacement, fast / slow time constant, slow time constant, torque range | control (6-8); injury 1 (5-8); injury 2 (4-8) | compression-tension, compressive creep, axial rotation |
| 22 | Norcross (2003)<sup>56</sup> | rat (male) | unknown (unknown) | tail | biochemical | CABC | 0.06-0.1mL (0.25U/mL) | AF+NP | 2 | stiffness, hysteresis, residual deformation | control (20); injury (20) | compression-tension, compressive creep |
| 23 | Rousseau (2007)<sup>24</sup> | rat (unknown) | 3 month (unknown) | lumbar, tail | physical disruption | stab lesion | no.11 blade, 1.5mm depth | AF+NP | 4 | stiffness, NZ RoM | lumbar: control (10), injury (12); tail: control (11), injury (9) | flexion-extension, lateral bending |
| 24 | Wakano (1983)<sup>35</sup> | dog (unknown) | unknown (25 kg) | lumbar | biochemical | chymopapain | 0.1mL | NP | 3, 12 | stiffness, creep rate | control (intact:9, sham: 1); injury 1 (13); injury 2 (2) | compression-tension, axial rotation, compressive creep, torsional creep |
| 2<br>5 | Wei (2021)<sup>39</sup> | mouse<br>(female) | 10 week<br>(unknown) | tail | physical<br>disruption | needle<br>puncture | 26G | AF+NP | 4 | modulus,<br>hysteresis,<br>compressive<br>strain | control (4);<br>injury (4) | compression-tension,<br>compressive<br>creep |
| 2<br>6 | Xu (2024)<sup>57</sup> | rat (male) | unknown<br>(300 gram) | tail | physical<br>disruption | scalpel<br>rupture | no.11 blade,<br>depth: full AF<br>thickness | AF | 1, 4 | 3-point bending<br>force | control (3);<br>injury (3) | 3-point<br>bending |
| 2<br>7 | Yuan<br>(2022)<sup>36</sup> | goat<br>(female) | 24 month<br>(unknown) | lumbar | physical<br>disruption | discectomy +<br>AF suture | 50 mg NP tissues<br>was removed | AF+NP | 24 | RoM | control<br>(10);<br>injury (10) | flexion-extension,<br>lateral<br>bending, axial<br>rotation |
| 2<br>8 | Detiger<br>(2013)<sup>38</sup> | goat<br>(female ) | 3-4 year<br>(92 ± 10 kg) | lumbar | biochemical | CABC | 130μL (0.25U/mL | NP | 12 | NZ stiffness,<br>RoM, NP elastic<br>/ viscous<br>modulus | control<br>(14-16);<br>injury (14-16) | flexion-extension,<br>lateral<br>bending, axial<br>rotation,<br>intradiscal |

*IVD, intervertebral disc; IVDD, intervertebral disc degeneration; CABC, chondroitinase ABC; AF, annulus fibrosis; NP, nuclear pulposus; NZ, neutral zone; RoM, range of motion*

**Table 3.** Characteristics of Included Studies Reporting Morphological Outcomes.

| ID | Author (Year) | Spine Level | Assessment Method | Timepoints (weeks) | Reported Outcomes (with Scale) | Groups (n) |
| --- | --- | --- | --- | --- | --- | --- |
| 1 | Boxberger (2008)<sup>55</sup> | lumbar | μCT | 4, 12 | DH | control (13); injury (13) |
| 3 | Calvo-Echenique (2018)<sup>51</sup> | lumbar | MRI | 12, 24 | DH | control (4); injury (8) |
| 4 | Colloca (2012)<sup>40</sup> | lumbar | Histology (H&E) | 6, 13 | degeneration grade (1-4) | control 1 (6); injury 1 (6) |
| 5 | Crowley (2024)<sup>49</sup> | lumbar | X-ray, MRI, Histology (H&E) | 6, 12 | DHI, degeneration grade (Pfirrmann: 1-5; histology: 0-14) | control 1 (6); injury 1 (6); control 2 (6); injury 2 (6) |
| 6 | Fainor (2023)<sup>50</sup> | lumbar | Histology (H&E, Safranin-O/Fast Green) | 4, 10 | degeneration grade (scale not reported) | control (4-6); injury (4-6) |
| 7 | Fazzalari (2001)<sup>41</sup> | lumbar | Macroscopic grading (visual inspection) | 0, 4, 12, 24, 48, 72 | degeneration grade (1-4) | control (4); injury (48) |
| 8 | Gullbrand (2017)<sup>37</sup> | lumbar | MRI, Histology (H&E, AB&PR) | 12 | DHI, degeneration grade (Pfirrmann: 1-5; histology: 0-100) | control (5-10); injury (10) |
| 9 | He (2022)<sup>53</sup> | tail | X-ray, Histology (H&E, Safranin O, Alcian Blue) | 2 | degeneration grade (4-12) | control (10); injury (10) |
| 13 | Latham (1994)<sup>42</sup> | lumbar | Histology (H&E) | 24 | degeneration category (none, mild, moderate, severe) | control (6); injury (12) |
| 14 | Leckie (2012a)<sup>44</sup> | lumbar | MRI | 4 | degeneration grade (Pfirrmann: 1-5) | control (6); injury (2-4) |
| 17 | Lü (1997)<sup>34</sup> | lumbar | X-ray | 1 | DHI | control (8); injury 1 (16); injury 2 (16); injury 3 (8) |
| 18 | Martin (2013)<sup>21</sup> | tail | μCT | 8 | DH | control (7); injury 1 (7); injury 2 (7) |
| 19 | Melrose (2012)<sup>23</sup> | lumbar | MRI | 12 | DH | control (7); injury (6) |
| 20 | Mosley (2020)<sup>25</sup> | lumbar | X-ray, Histology (AB&PR) | 6 | DH, degeneration grade (histology: 0-10) | control (7-12); injury (7-12) |
| 21 | Mosley (2019)<sup>54</sup> | lumbar | X-ray, Histology (AB&PR) | 6 | DH, degeneration grade (histology: 0-10) | control (6-8); injury (6-8) |
| 22 | Norcross (2003)<sup>56</sup> | tail | X-ray, Histology (Alcian Blue) | 2 | DH, degeneration grade (histology: 1-5) | control (10-20); injury (10-20) |
| 26 | Xu (2024)<sup>57</sup> | tail | X-ray, Histology (H&E) | 1, 4 | DH, degeneration grade (histology: 5-15) | control 1 (20); injury 1 (20); control 2 (3); injury 2 (3) |
| 27 | Yuan (2022)<sup>36</sup> | lumbar | X-ray, MRI | 24 | DHI, degeneration grade (pfirrmann: 1-5, histology: 0-5) | control (9-10); injury (9-10) |
| 28 | Detiger (2013)<sup>38</sup> | lumbar | X-ray | 12 | DHI | control (24); injury (24) |

*Only studies reporting disc height (DH) or degeneration grade were included. Studies reporting categorical outcomes were not included in quantitative synthesis. DH, disc height; DHI, disc height index; H&E, hematoxylin and eosin; AB&PR, Alcian Blue and Picrosirius Red staining*

Seven species were represented: dog^34^ ^35^, goat^36–38^, mouse^21^ ^39^, ovine^23^ ^40–42^, porcine^43–48^, rabbit^49–52^, and rat^24^ ^25^ ^53–57^. Eleven studies^23^ ^34^ ^37^ ^41^ ^42^ ^44^ ^50^ ^53^ ^55–57^ used only male animals, five^36^ ^38^ ^39^ ^49^ ^52^ used only females, two^25^ ^54^ used both sexes, and ten^21^ ^24^ ^35^ ^40^ ^43^ ^45–48^ ^51^ did not report sex. Lumbar IVDs were most commonly used (82%, 23 of 28 studies) ^23–25^ ^34–38^ ^40–52^ ^54^ ^55^, with tail IVDs used in six studies^21^ ^24^ ^39^ ^53^ ^56^ ^57^. Three IVD injury models were identified: physical disruption (e.g., needle puncture, stab lesion, nucleotomy, fenestration) ^21^ ^23^ ^24^ ^36^ ^39–43^ ^45–53^ ^57^, biochemical (e.g., chondroitinase ABC, chymopapain, TNF-alpha injection) ^34^ ^35^ ^38^ ^44^ ^55^ ^56^, and combined physical plus biochemical disruption^25^ ^54^. Timepoints of mechanical testing following IVD injury ranged from 0 to 72 weeks.

### Methodological Quality Assessment

The methodological quality assessment of the 28 included studies is presented in Table 4. The median quality score was 7.3 (IQR 1.5), with scores ranging from 4 to 9.5. All studies were published in peer-reviewed journals. Blinding of outcome assessment was reported in only nine studies. Sample size calculations were documented in just three studies. While all studies clearly described their control groups, some used less appropriate controls, such as adjacent IVD(s) from the same animal. Statement of compliance with regulatory requirements were presented in all but one study^35^. Key methodological details, including the IVDD animal model, timing of outcome assessment, and primary and secondary outcomes were clearly reported across all studies. Statistical methods were appropriately described and justified in 93% of studies. A statement regarding conflicts of interest was provided by 19 out of 28 studies (68%).

**Table 4.** Quality of included studies based on the CAMARADES checklist.

| Study | 1 | 2 | 3 | 4 | 5 | 6 | 7 | 8 | 9 | 10 | Quality score |
| --- | --- | --- | --- | --- | --- | --- | --- | --- | --- | --- | --- |
| Boxberger (2008) | 1 | 0.5 | 0 | 0.5 | 1 | 1 | 1 | 1 | 1 | 1 | 8 |
| Buser (2011) | 1 | 0.5 | 0 | 0.5 | 1 | 1 | 1 | 0.5 | 1 | 0 | 6.5 |
| Calvo-Echenique (2018) | 1 | 0 | 0 | 1 | 1 | 1 | 1 | 0.5 | 1 | 1 | 7.5 |
| Colloca (2012) | 1 | 0 | 0 | 1 | 1 | 0.5 | 1 | 1 | 1 | 1 | 7.5 |
| Crowley (2024) | 1 | 1 | 1 | 1 | 1 | 1 | 1 | 0.5 | 1 | 1 | 9.5 |
| Fainor (2023) | 1 | 0.5 | 0 | 1 | 1 | 0.5 | 1 | 1 | 1 | 1 | 8 |
| Fazzalari (2001) | 1 | 0 | 0 | 0.5 | 0.5 | 0.5 | 1 | 1 | 1 | 0.5 | 6 |
| Gullbrand (2017) | 1 | 0.5 | 0 | 0.5 | 1 | 1 | 1 | 1 | 1 | 1 | 8 |
| He (2022) | 1 | 0 | 0 | 1 | 1 | 0.5 | 1 | 1 | 1 | 1 | 7.5 |
| Kaigle (1997) | 1 | 0 | 0 | 1 | 0 | 1 | 1 | 0.5 | 1 | 0 | 5.5 |
| Kaigle (1998) | 1 | 0 | 0 | 1 | 0 | 0.5 | 1 | 1 | 1 | 0 | 5.5 |
| Keller (1990) | 1 | 0 | 0 | 0.5 | 0 | 0.5 | 1 | 1 | 1 | 0 | 5 |
| Latham (1994) | 1 | 0 | 0.5 | 1 | 1 | 1 | 1 | 1 | 0.5 | 0 | 7 |
| Leckie (2012a) | 1 | 0 | 1 | 1 | 1 | 0.5 | 1 | 1 | 1 | 0.5 | 8 |
| Leckie (2012b) | 1 | 0 | 1 | 1 | 1 | 0.5 | 1 | 1 | 0 | 1 | 7.5 |
| Lin (2015) | 1 | 0.5 | 0 | 0.5 | 1 | 1 | 1 | 0.5 | 1 | 0 | 6.5 |
| Lü (1997) | 1 | 0 | 0 | 0.5 | 1 | 0.5 | 1 | 0.5 | 0.5 | 0 | 5 |
| Martin (2013) | 1 | 0 | 0 | 0.5 | 1 | 0.5 | 1 | 1 | 1 | 0.5 | 6.5 |
| Melrose (2012) | 1 | 0 | 0 | 1 | 1 | 0.5 | 1 | 0.5 | 1 | 1 | 7 |
| Mosley (2020) | 1 | 0.5 | 0 | 1 | 1 | 0.5 | 1 | 1 | 1 | 1 | 8 |
| Mosley (2019) | 1 | 0.5 | 0 | 1 | 1 | 1 | 1 | 1 | 1 | 0.5 | 8 |
| Norcross (2003) | 1 | 0.5 | 0 | 0.5 | 1 | 0.5 | 1 | 1 | 1 | 0 | 6.5 |
| Rousseau (2007) | 1 | 0 | 0 | 1 | 1 | 0.5 | 1 | 1 | 1 | 1 | 7.5 |
| Wakano (1983) | 1 | 0 | 0 | 0.5 | 0 | 0.5 | 1 | 1 | 0 | 0 | 4 |
| Wei (2021) | 1 | 0 | 0 | 1 | 1 | 1 | 1 | 1 | 1 | 1 | 8 |
| Xu (2024) | 1 | 0 | 0 | 1 | 1 | 1 | 1 | 0.5 | 1 | 0.5 | 7 |
| Yuan (2022) | 1 | 0.5 | 0 | 0.5 | 1 | 1 | 1 | 0.5 | 1 | 1 | 7.5 |
| Detiger (2013) | 1 | 0 | 0 | 0.5 | 1 | 0.5 | 1 | 1 | 1 | 1 | 7 |

*Studies fulfilling the criteria of (1) peer-reviewed publication; (2) blinding of outcome assessment; (3) sample size calculation; (4) control group; (5) animal welfare; (6) animal model; (7) timing of outcome assessment; (8) primary and secondary outcomes; (9) statistical analysis; and (10) conflict of interests.*

### Risk of Bias in Included Studies

The risk of bias in the 28 included studies was assessed using the SYRCLE tool and is summarized in Figure 2. Regarding selection bias, 68% of the studies had low risk based on baseline characteristics. However, most studies presented an unclear risk of bias for sequence generation and allocation concealment, with the exception of one study^35^, which was rated as high risk for sequence generation. Performance bias was generally unclear across all studies due to a lack of reporting on random housing. Blinding of investigators was unclear in most studies. Only two studies had low risk, five studies had high risk. For detection bias, the majority of studies presented unclear risk, with only one study^41^ having high risk of random outcome assessment, and three studies having high risk of outcome assessor blinding. Attrition bias was low in most studies (79%) with adequate reporting of incomplete outcomes, although several studies presented a high risk. Reporting bias was generally low risk in 86% of studies, while four studies showed high risk due to incomplete reporting of pre-specified outcomes or selectively reporting of p-values without accompanying numerical or graphical data. Most studies were free from other sources of bias. However, a few were rated high risk primarily because they used adjacent IVD(s) from the same animal as controls.

**Figure 2.**
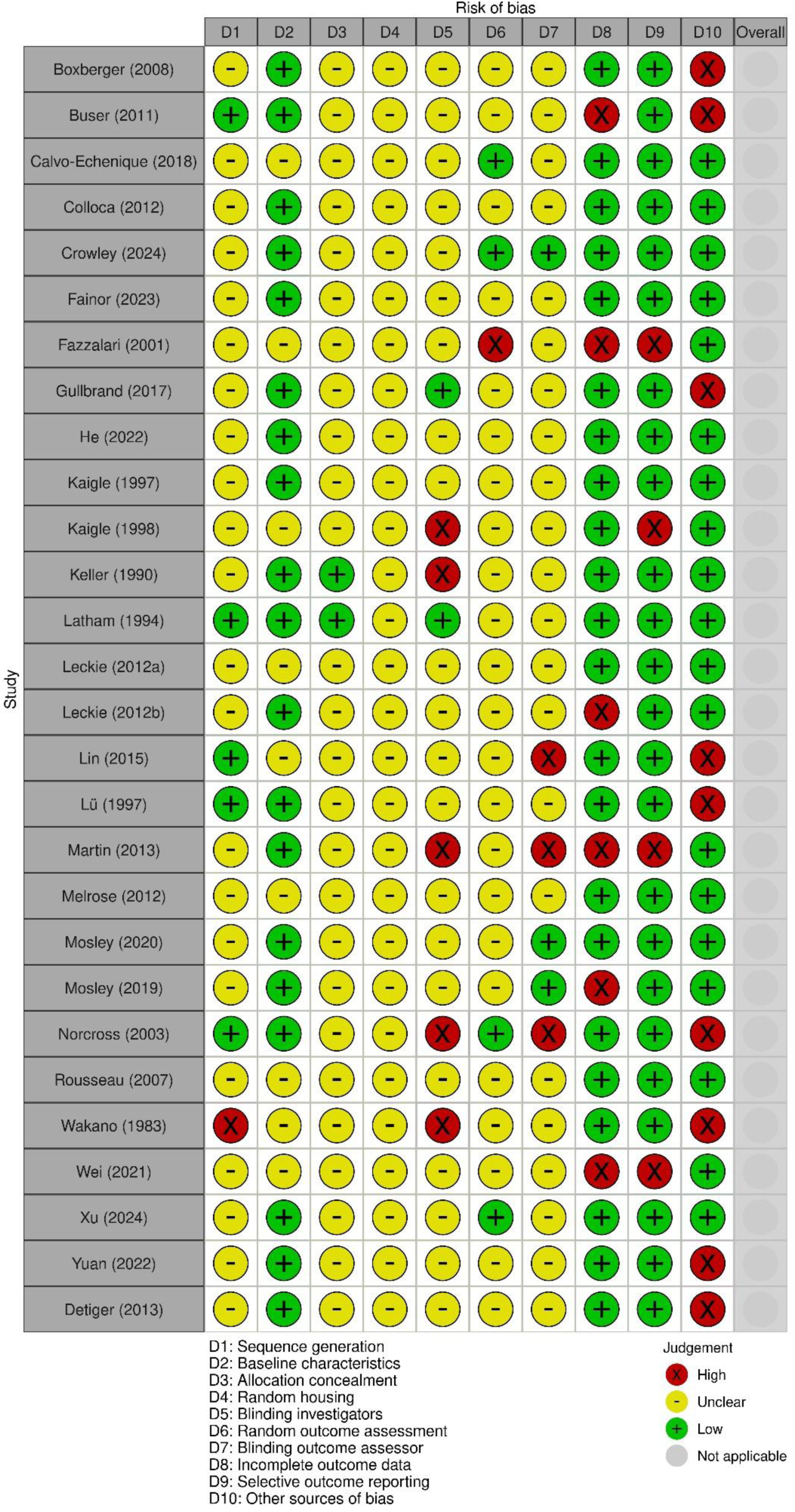
SYRCLE risk of bias assessment summary

### Effect sizes

#### Stiffness

55 effect sizes from eight studies^24^ ^25^ ^35^ ^38^ ^42^ ^44^ ^52^ ^54^ were initially included, one effect size^25^ was excluded based on sensitivity analysis. Meta-analysis of the remaining 54 effect sizes showed a significant reduction in stiffness following IVD injury (SMD=-0.24, SE=0.10, 95% CI: −0.42 to −0.05, p=0.013) (Figure 3A), with moderate heterogeneity (τ²=0.16, I²=34.22%, Q(53)=93.29, p<0.001). The prediction interval (−1.05 to 0.57) indicated that future studies may observe a wider range of effects, including potentially null or even positive effects. Egger’s test indicated evidence of publication bias (z=-2.14, p=0.033). Funnel plots for all outcome domains are provided in the Supplementary figure (Figure S1).

**Figure 3.**
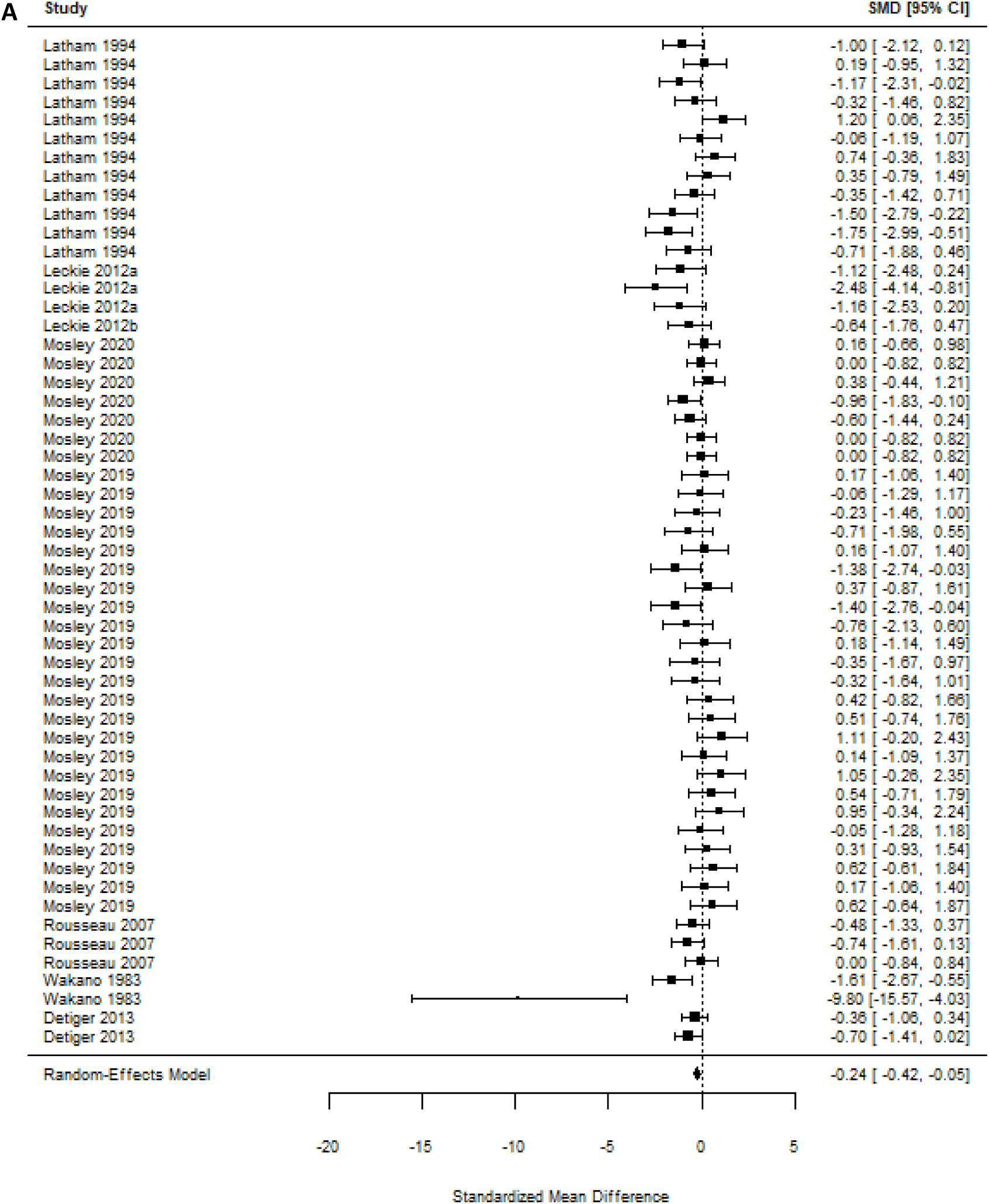
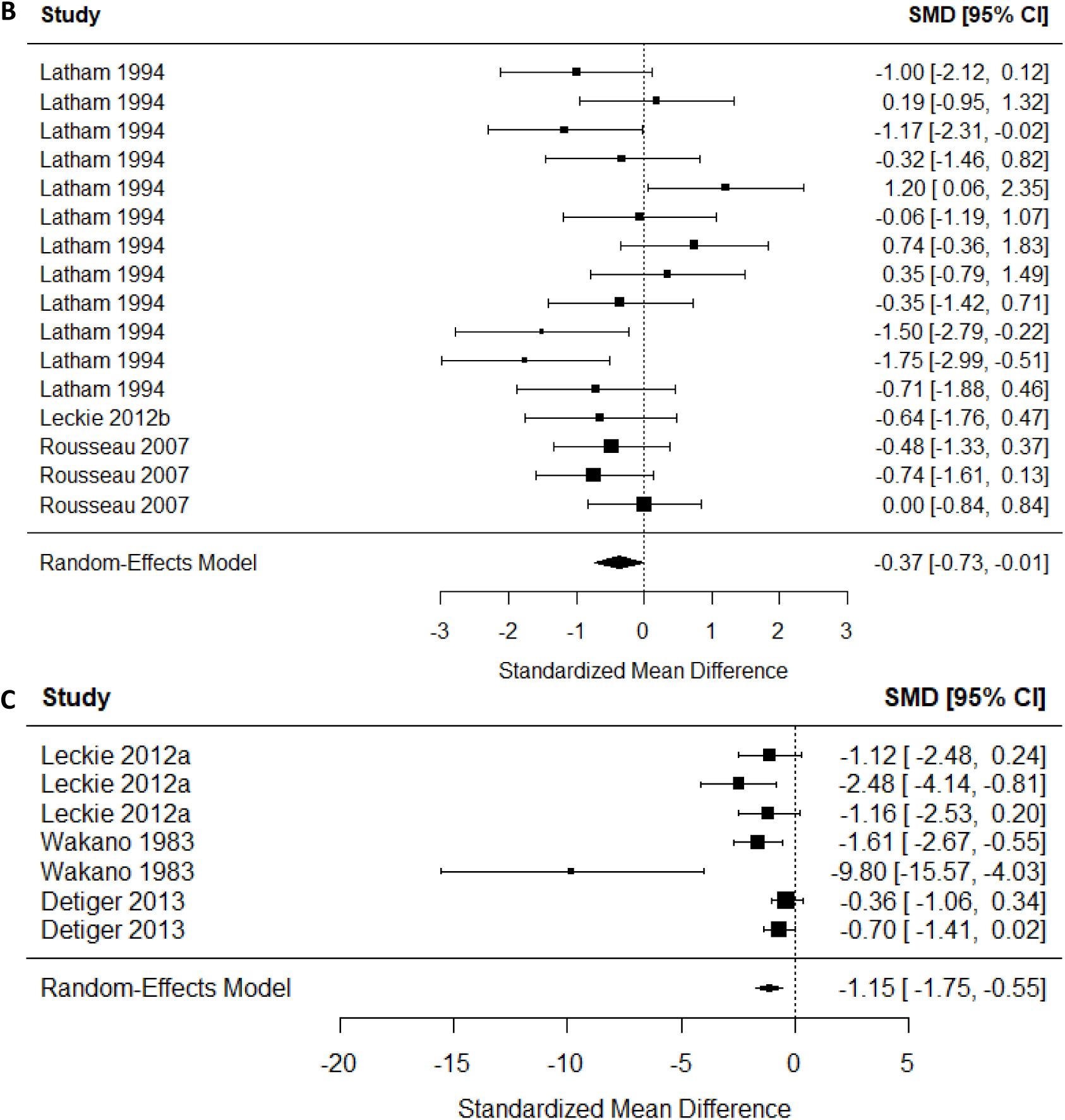
Forest plots of the Standardized Mean Difference estimates, 95% confidence interval (CI), and prediction intervals of IVD injury effects on changes in stiffness for (A) all models, (B) physical disruption models, and (C) biochemical models.

Univariate moderator analyses identified species, injury model, damaged structure, outcome parameters, and publication year as significant moderators. However, species was excluded from meta-regression due to sparse representation across studies. In the final meta-regression model, injury model emerged as a significant source of heterogeneity (QM(5)=16.38, p=0.006). Subgroup analysis of physical disruption models (k=16, 3 studies) ^24^ ^42^ ^52^ showed a significant reduction in stiffness following IVD injury (SMD=-0.37, SE=0.18, 95% CI: −0.73 to −0.01, p=0.042) (Figure 3B), with moderate heterogeneity (τ²=0.23, I²=43.14%, Q(15)=26.80, p=0.030) and no evidence of publication bias (z=-0.67, p=0.50). Subgroup analysis of biochemical models (k= 7, 3 studies) ^35^ ^38^ ^44^ also showed a significant reduction in stiffness following IVD injury (SMD=-1.15, SE=0.31, 95% CI: −1.75 to −0.55, p<0.001) (Figure 3C), with moderate heterogeneity (τ²=0.26, I²=45.12%, Q(6)=17.07, p=0.009), but with evidence of publication bias (z=-3.84, p<0.001). No further subgroup analysis was conducted for physical plus biochemical disruption models (k=31, two studies) ^25^ ^54^ due to limited studies.

#### Young’s Modulus

Although species was a significant moderator in univariable analysis (QM(3)=14.09, p=0.003), each species level was represented by a single study. Thus, meta-regression was not pursued, and a pooled analysis across species was conducted. Random-effects analysis of 18 effect sizes from four studies^21^ ^35^ ^37^ ^55^ showed a significant reduction in Young’s modulus following IVD injury (SMD=-1.22, SE=0.22, 95% CI: −1.64 to −0.80, p<0.001) (Figure 4A), with moderate heterogeneity (τ^2^=0.39, I^2^=52.95%, Q(17)=34.46, p=0.007). The prediction interval ranged from −2.52 to 0.08. No influential studies were detected, but Egger’s test indicated significant publication bias (z=-3.63, p<0.001).

**Figure 4.**
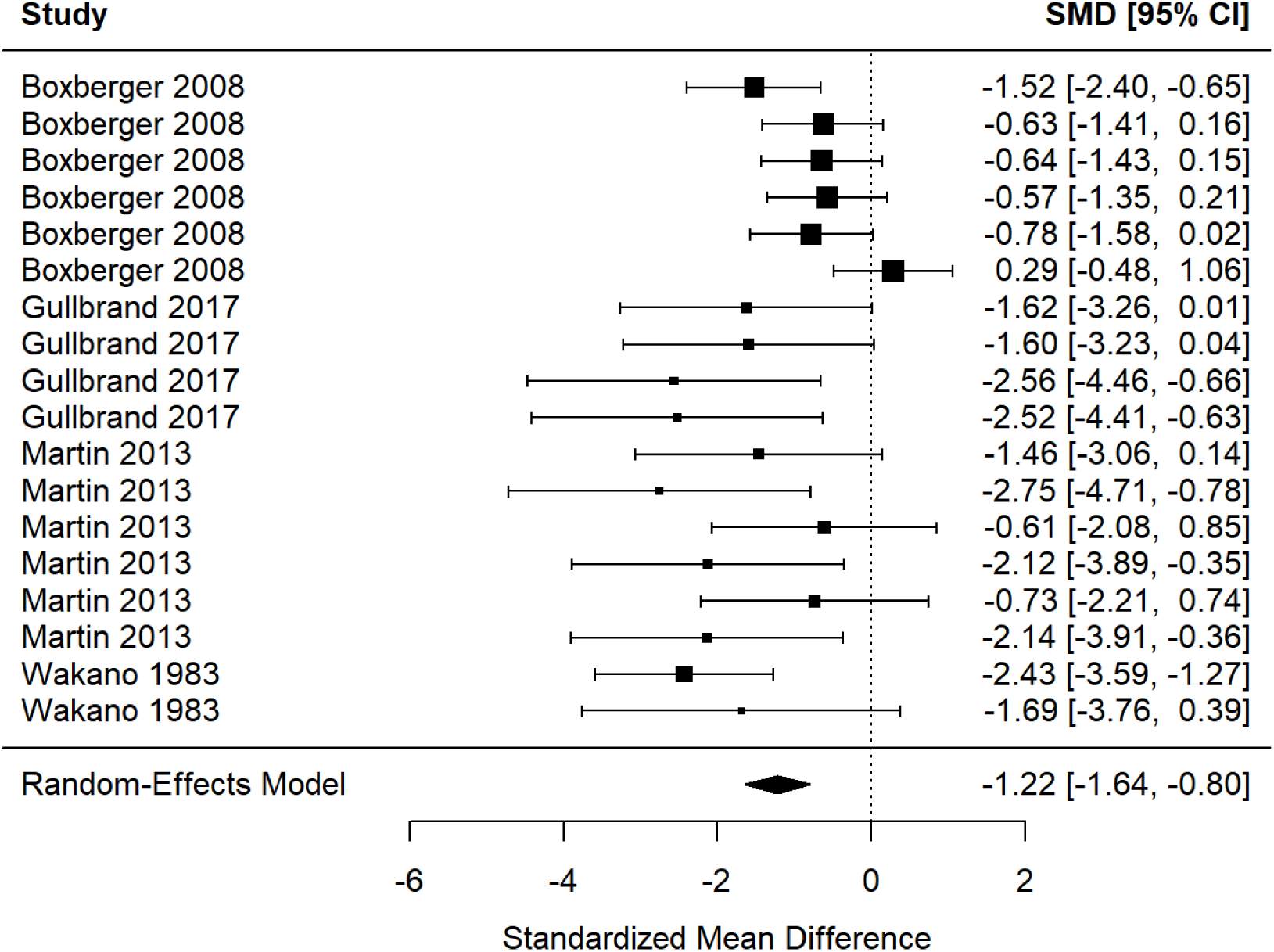
Forest plots of the Standardized Mean Difference estimates, 95% confidence interval (CI), and prediction intervals of IVD injury effects on changes in Young’s modulus.

#### RoM

79 effect sizes from 13 studies^21^ ^23–25^ ^34^ ^36–38^ ^44^ ^49^ ^54^ ^55^ were initially included. One influential effect size^36^ was excluded after sensitivity analysis. Meta-analysis of the remaining 78 effect sizes showed a significant increase in RoM following IVD injury (SMD=0.74, SE=0.17, 95% CI: 0.40 to 1.08, p<0.001) (Figure 5A), with substantial heterogeneity (τ²=1.86, I²=82%, Q(77)=335.05, p<0.001). The prediction interval (−1.95 to 3.43) indicated inconsistent direction of effects in future studies. Egger’s test indicated evidence of publication bias (z=2.91, p=0.004). Univariable moderator analyses identified species, sex, injury model, publication year, and timepoints as significant moderators. Species and publication year were excluded from meta-regression due to sparse representation across studies.

**Figure 5.**
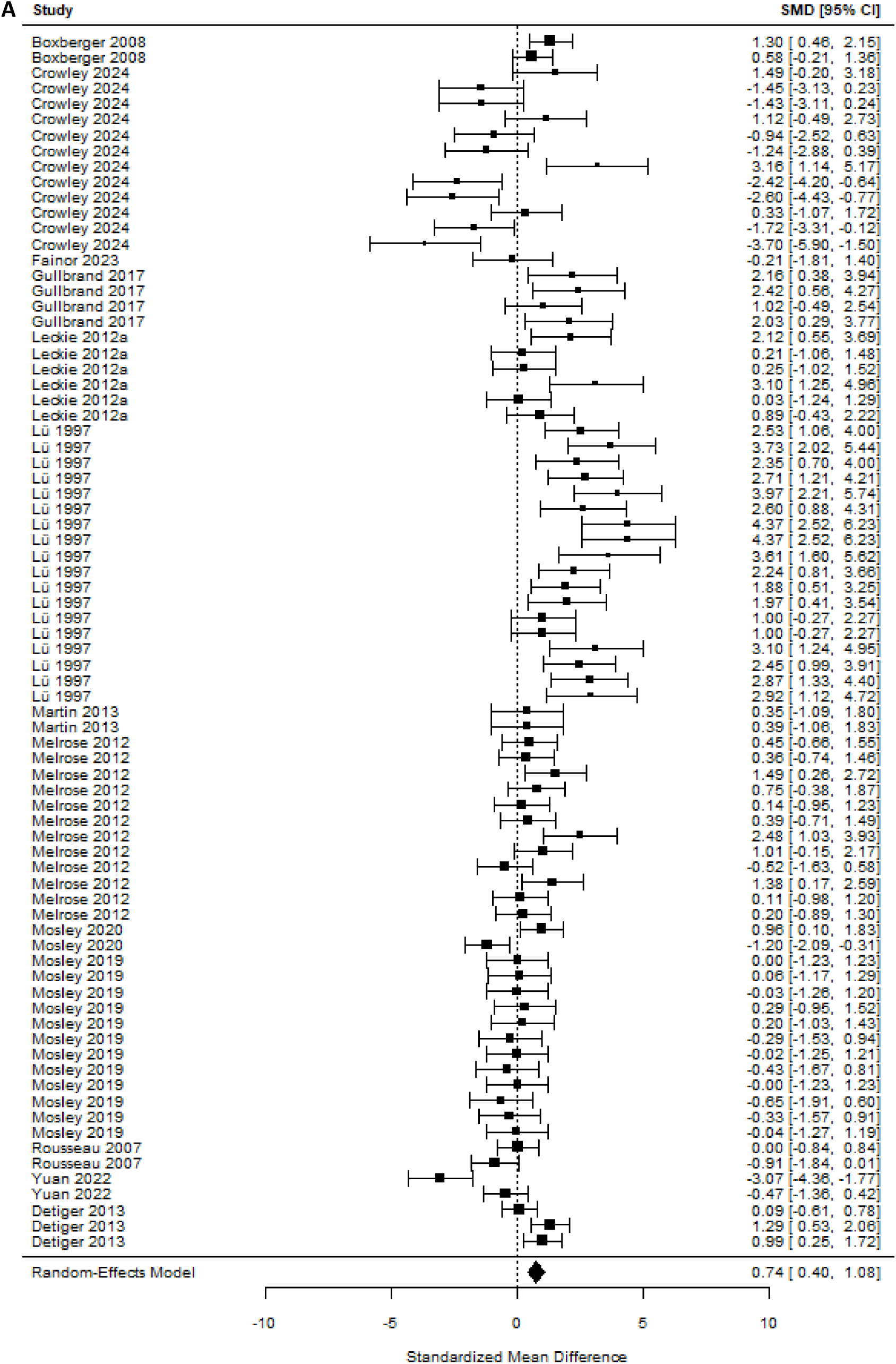
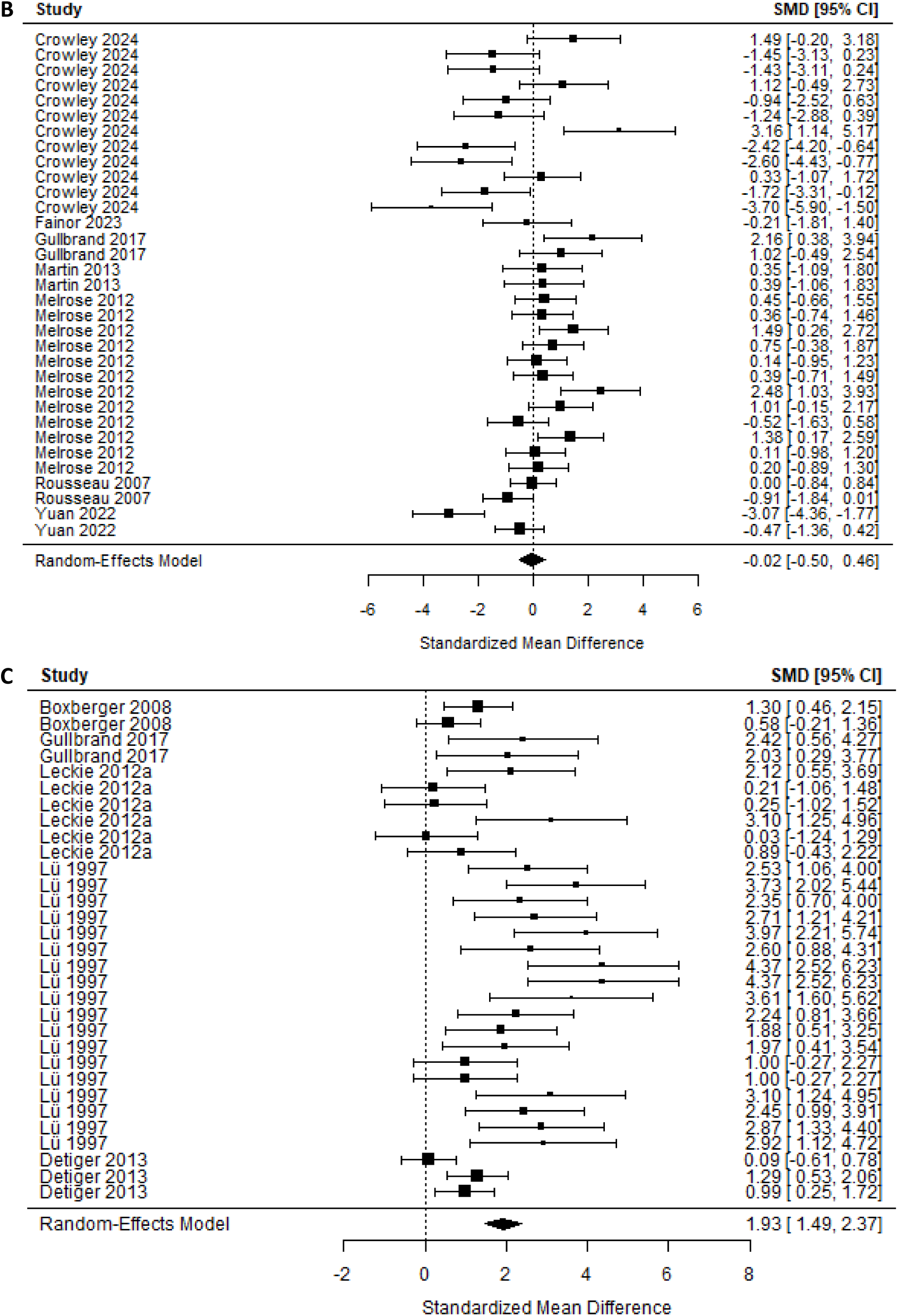
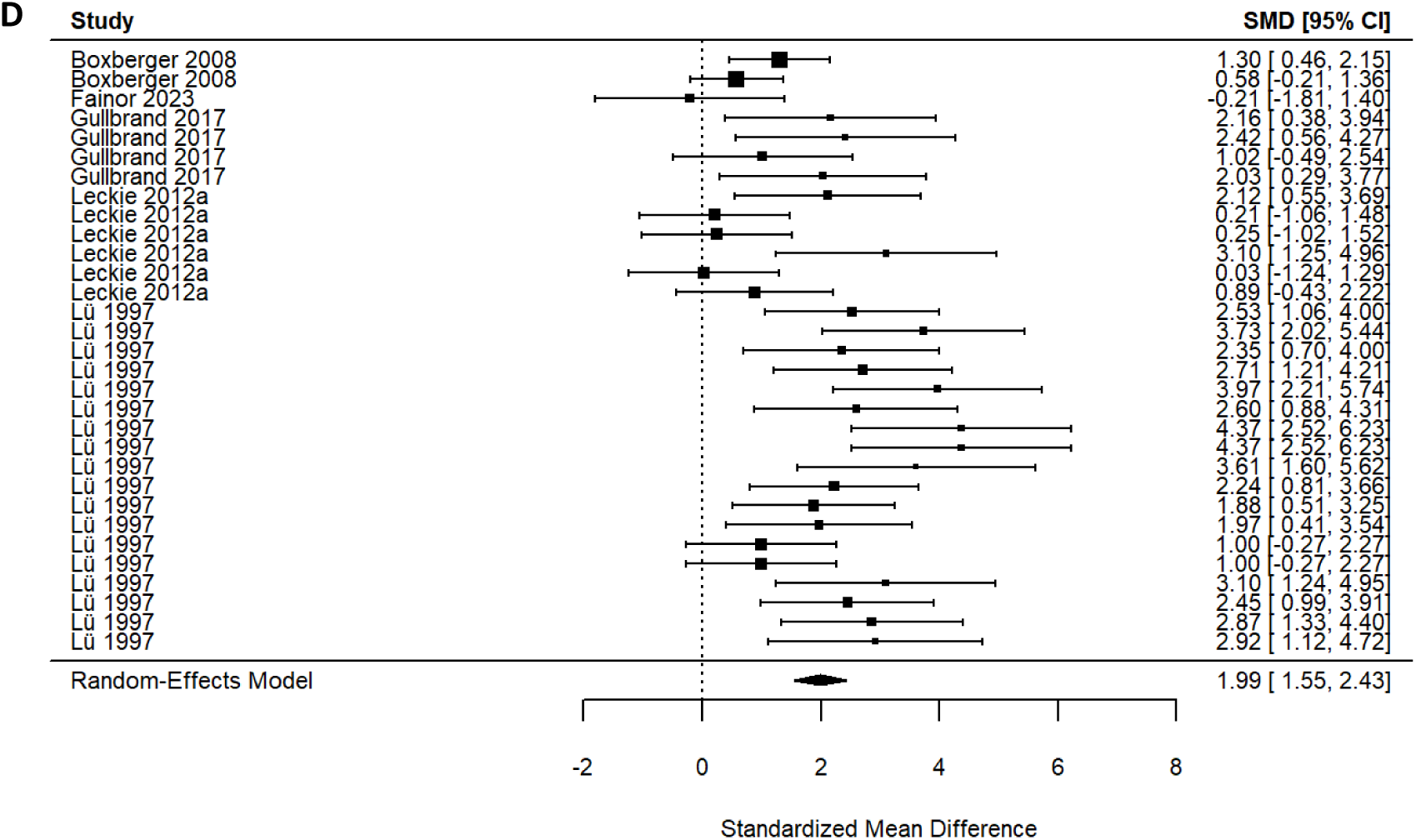
Forest plots of the Standardized Mean Difference estimates, 95% confidence interval (CI), and prediction intervals of IVD injury effects on changes in RoM (range of motion) for (**A**) all models, (**B**) physical disruption models, (**C**) biochemical models, and (**D**) male models.

In the final meta-regression model, injury model and sex emerged as significant sources of heterogeneity (QM(5)=79.06, p<0.001). Subgroup analysis of physical disruption models (k=33, 7 studies) ^21^ ^23^ ^24^ ^36^ ^37^ ^49^ ^50^ showed a non-significant reduction in RoM following IVD injury (SMD=-0.02, SE=-0.10, 95% CI: −0.50 to 0.46, p=0.923) (Figure 5B), with substantial heterogeneity (τ²=1.47, I²=77.18%, Q(32)=117.40, p<0.001) and a prediction interval ranging from −2.45 to 2.40. No evidence of publication bias was detected (z=-10, p=0.319). Subgroup analysis of biochemical models (k= 31, 5 studies) ^34^ ^37^ ^38^ ^44^ ^55^ showed a significant increase in RoM following IVD injury (SMD=1.93, SE=0.23, 95% CI: 1.49 to 2.37, p<0.001) (Figure 5C), with moderate heterogeneity (τ²=1.03, I²=71.10%, Q(30)=98.52, p<0.001), but with evidence of publication bias (z=6.35, p<0.001). The prediction interval ranged from −0.11 to 3.96. No further subgroup analysis was conducted for combined physical disruption plus biochemical models (k=14, two studies) ^25^ ^54^ due to limited studies.

Subgroup analysis of male models (k=31, 5 studies) ^34^ ^37^ ^44^ ^50^ ^55^ showed a significant increase in RoM following IVD injury (SMD=1.99, SE=0.23, 95% CI: 1.55 to 2.43, p<0.001) (Figure 5D), with moderate heterogeneity (τ²=0.97, I²=64.39%, Q(30)=82.75, p<0.001), prediction interval ranged from 0.01 to 3.97, but evidence of publication bias was detected (z=5.05, p<0.001). Subgroup analysis of female models (k= 17, 3 studies) showed a non-significant decrease in RoM following IVD injury (SMD=-0.57, SE=0.43, 95% CI: −1.40 to 0.27, p=0.184), with substantial heterogeneity (τ²=2.51, I²=86.37%, Q(16)=94.28, p<0.001), there was no evidence of publication bias (z=-1.43, p=0.154). The prediction interval (−3.78 to 2.65) supported an inconsistent direction of effect.

#### viscoelasticity

Eighty-one effect sizes from nine studies^21^ ^25^ ^35^ ^37^ ^50^ ^51^ ^54–56^ were included in the analysis. The random-effects model showed no significant change in viscoelastic properties of the IVD following injury (SMD=0.01, SE=0.13, 95% CI: −0.25 to 0.26, p=0.953) (Figure 6). Substantial heterogeneity was observed across studies (τ²=0.91, I²=71.36%, Q(80)=269.59, p<0.001), with a prediction interval ranging from −1.88 to 1.90. Because moderator levels were sparsely represented across studies, meta-regression was not performed. No influential studies were detected in sensitivity analysis, and Egger’s test indicated no evidence of publication bias (z=0.598, p=0.550).

**Figure 6.**
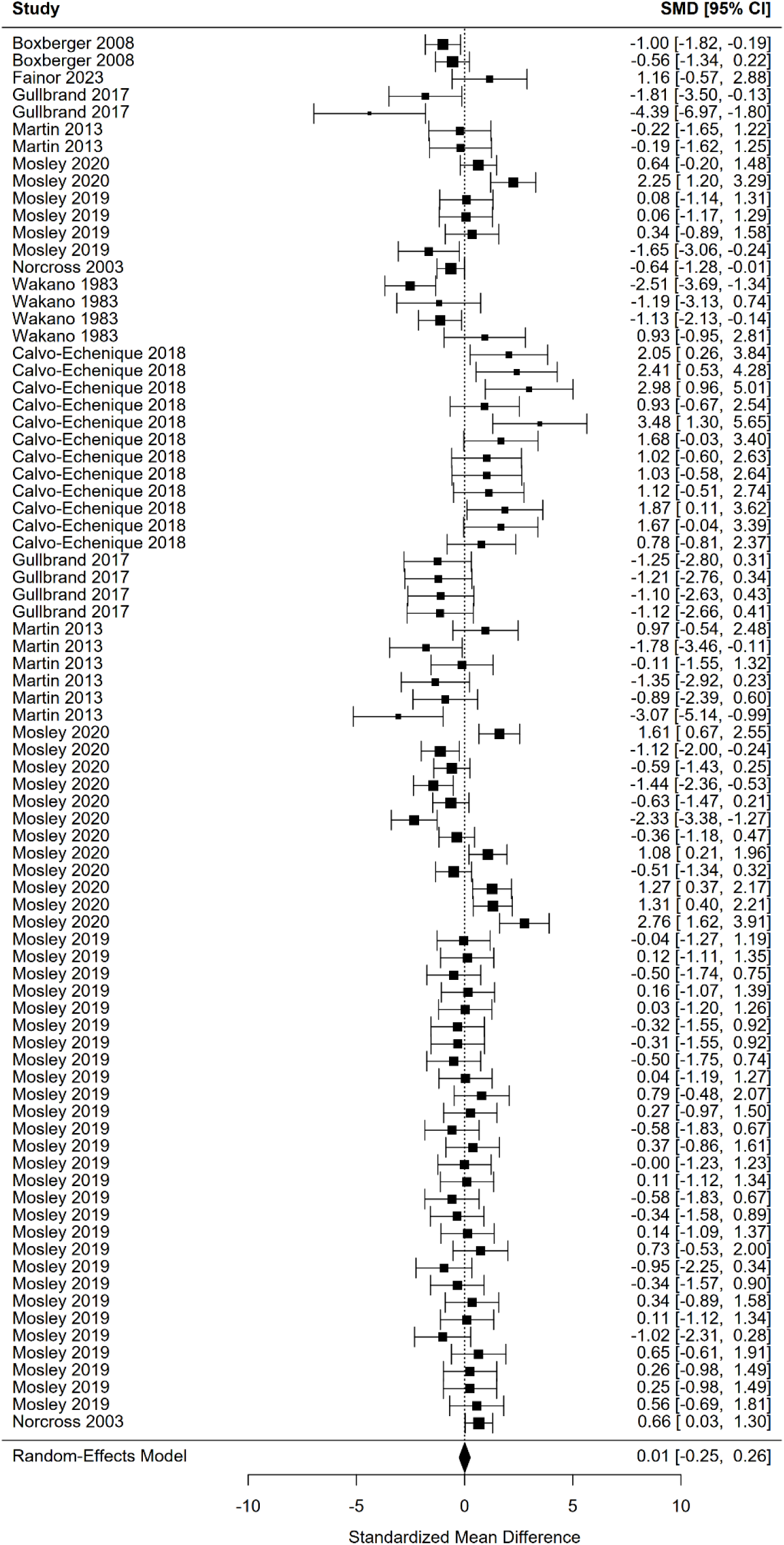
Forest plots of the Standardized Mean Difference estimates, 95% confidence interval (CI), and prediction intervals of IVD injury effects on changes in viscoelasticity.

#### Pressure-leakage

Eight records from one single study were available for this domain^43^. Thus, no meta-analysis or moderator analysis was conducted, and findings are presented descriptively. In this study, IVD injury was associated with decreased leakage and saturation pressure (SMD range: −4.11 to −4.98) and decreased leakage volume (SMD range: −1.38 to −1.68), while saturation volume showed inconsistent changes (SMD range: −0.95 to 0.69). Given that these findings are derived from a single study, they should be interpreted with caution.

#### Disc height

26 effect sizes from 12 studies^21^ ^25^ ^34^ ^37^ ^38^ ^49^ ^51^ ^53–57^ were initially included. One influential effect size^25^ was excluded after sensitivity analysis. Univariable moderator analyses identified species and timepoint as significant moderators. Multivariable meta-regression (k=25, 12 studies) including these two moderators was statistically significant (QM(6)=71.33, p<0.001), explaining 94.64% of the heterogeneity (R²=94.64%, τ²=0.20, I²=32.84%).

Based on the meta-regression results and the distribution of available studies, subgroup meta-analysis was performed for the rat species only, which had sufficient data from six studies^25^ ^53–57^, while other species (dog, goat, mouse, ovine, rabbit) were represented by one or two studies. The pooled effect size of rat was calculated (k=11), and it showed a significant decrease of disc height (SMD=-2.17, SE=0.63, 95% CI: −3.41 to −0.93, p<0.001). Two influential studies^25^ ^53^ were excluded after sensitivity analysis, which did not alter the significance of the injury effects (SMD=-1.29, SE=0.21, 95% CI: −1.70 to −0.87, p<0.001) (Figure 7A). The prediction interval ranged from −1.70 to −0.86. Heterogeneity was substantial (τ²=1.14, I²=78.17%, Q(9)=30.68, p<0.001), and Egger’s test indicated no evidence of publication bias (z=-0.08, p=0.935).

**Figure 7.**
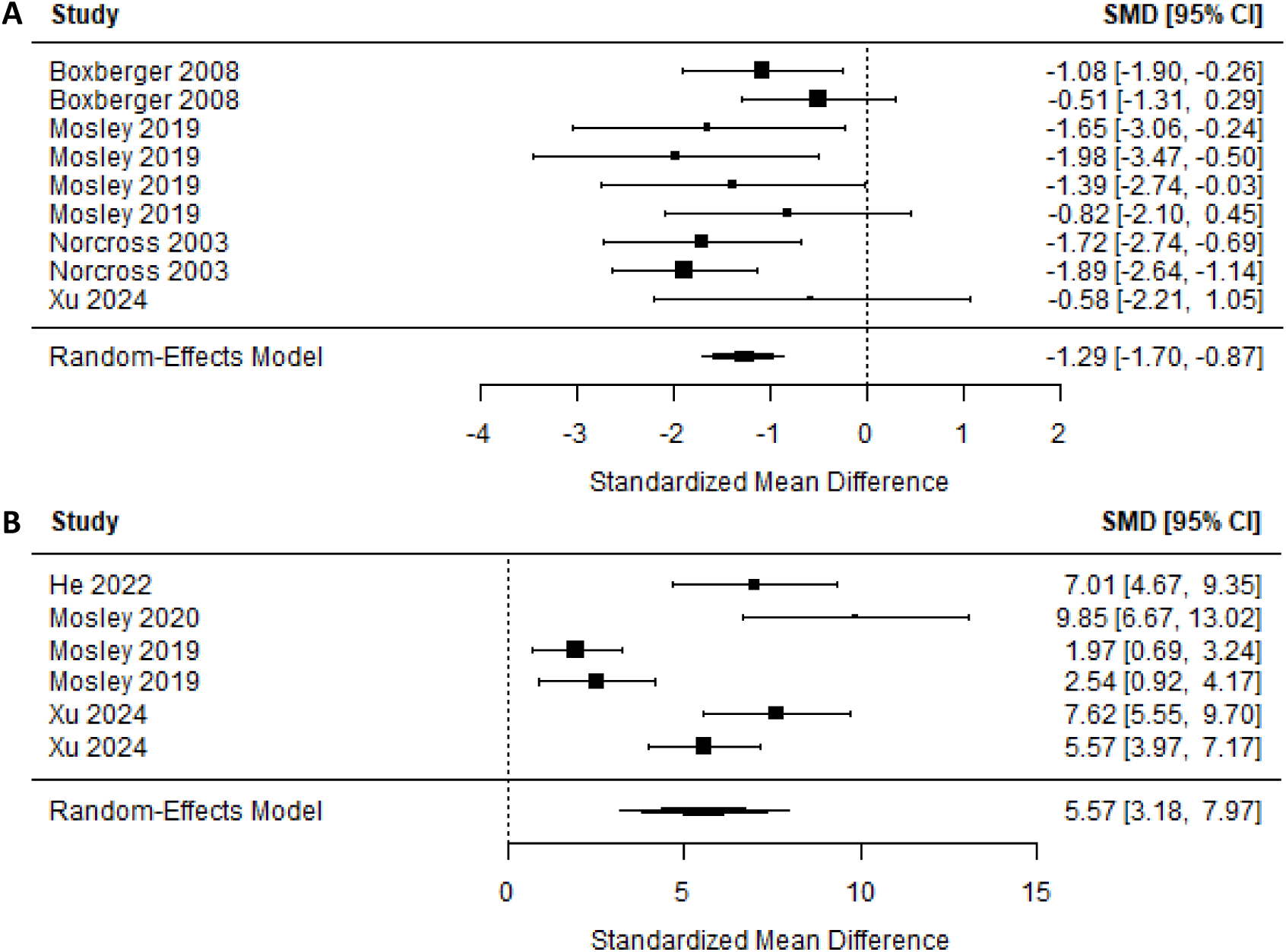
Forest plots of the Standardized Mean Difference estimates, 95% confidence interval (CI), and prediction intervals of IVD injury effects on changes in (**A**) disc height and (**B**) degeneration grade.

#### Degeneration grade

14 effect sizes from seven studies^25^ ^37^ ^40^ ^49^ ^53^ ^54^ ^57^ were initially included. Univariable moderator analyses identified species and sex as significant moderators. A multivariable meta-regression including these two moderators was statistically significant (QM(4)=22.70, p<0.001), explaining 67.78% of the heterogeneity (R²=67.78%, τ²=8.68, I²=87.26%).

Based on the meta-regression results and the distribution of available studies, subgroup meta-analysis was performed for the male rat subgroup, which was the only combination with sufficient data (k=6, four studies) ^25^ ^53^ ^54^ ^57^. The pooled effect size showed a significant increase in histological degeneration (SMD=5.57, SE=1.22, 95% CI: 3.18 to 7.97, p<0.001) (Figure 7B). No influential effect sizes were identified. Adding spine level (tail vs. lumbar) as a moderator did not significantly explain the heterogeneity (QM(1)=0.91, p=0.34), which remained substantial (τ²=7.58, I²=89.02%, Q(4)=23.23, p<0.001), and Egger’s test indicated evidence of publication bias (z=3.77, p<0.001). The prediction interval ranged from 3.18 to 7.97.

#### Relationship between morphological and mechanical outcomes

Some studies reported both morphological and mechanical outcomes following IVD injury, including measurements of disc height with stiffness^25^ ^38^ ^54^, Young’s modulus^21^ ^37^ ^55^, RoM^21^ ^25^ ^34^ ^37^ ^38^ ^49^ ^54^ ^55^, viscoelasticity^21^ ^25^ ^37^ ^51^ ^54–56^, as well as degeneration grade with stiffness^25^ ^54^, Young’s modulus^37^, RoM^25,37^ ^49^ ^54^ ^55^, viscoelasticity^25^ ^37^ ^49^ ^54^ ^55^. However, only a limited number of studies^25^ ^49^ ^54^ reported morphological and mechanical outcomes at the same IVD level. Due to the limited eligible studies with anatomically matched morphological and mechanical outcomes, formal correlation analysis was not feasible.

## Discussion

This systematic review and meta-analysis assessed the effects of IVD injury on mechanical properties and the relationship between morphological and mechanical changes in in vivo animal models of IVDD. To our knowledge, this is the first quantitative synthesis of mechanical changes across commonly used preclinical IVD injury models. Significant changes were observed in stiffness (reduction), Young’s modulus (reduction), range of motion (increase), disc height (reduction), and degeneration grade (increase), indicating a pattern of degeneration affecting both mechanical and structural domains. In contrast, viscoelasticity and creep-strain domains showed no non-significant changes, which may reflect methodological variability or sensitivity differences between outcome measures. Analysis of correlations between morphological and mechanical outcomes was not possible.

### Mechanical changes following IVD injury

A significant reduction in stiffness was observed following IVD injury, and subgroup analyses confirmed this reduction across different IVDD induction methods. However, the effect size was larger in biochemical models than in physical disruption models, possibly reflecting the more extensive disruption and depressurization of the NP in the biochemical models. Biochemical models use chemo-nucleolytic agents that degrade proteoglycans, aggregates, and glycosaminoglycan in the ECM, and these components are crucial to NP hydration and mechanical integrity^15^ ^19^. In contrast, physical disruption models may cause more localized damage, with injury severity varying according to anatomical location^58^, relative dimensions of instruments used^59^, number of punctions/stabs^60^, and penetration depth^61^. In both models, mechanical properties may evolve over time following injury^6^ ^62^ ^63^, and varying post-injury assessment timepoints have likely contributed to the variability in results.

A significant reduction in Young’s modulus was also observed following IVD injury. The injury effect was notably larger in Young’s modulus compared with stiffness, suggesting that modulus may be more sensitive to degenerative changes than stiffness. Although both parameters reflect disc mechanical function, they should not be used interchangeably, as they represent distinct aspects of tissue behavior. Young’s modulus is a geometry-independent measure of the disc’s material property. In contrast, stiffness is influenced by both material properties and geometry. For example, reductions in disc height have been shown to increase spinal stiffness^64^. Such geometric effects may partly explain why stiffness changes are less pronounced than modulus changes. Reporting both measures and specifying methods and disc dimensions is essential for comparability in future research.

RoM generally increased following IVD injury, but effect direction varied across studies. Heterogeneity in RoM changes was largely explained by model type and sex, with biochemical models and males showing more pronounced increases, though the observed sex effects may be partially cofounded by overlap between model type and sex distribution across studies^65^. Injury severity may further contribute to variability in RoM responses. Experimental evidence has shown that different injury models can produce distinct mechanical consequences. For instance, needle puncture has been reported to reduce peak stiffness in flexion, more than knife stab injury^66^. Such reductions in segmental stiffness would be expected to permit increased motion under load, which may contribute to the larger RoM observed in some experimental models^66^. Clinical evidence showed a non-linear relationship between degeneration severity and RoM in humans, segmental motion increased with degeneration severity Grade III to IV, then decreased at advanced degeneration grade V^67^ ^68^.

Viscoelastic properties did not demonstrate significant changes following IVD injury, likely due to methodological variability of testing protocols (e.g., dynamic testing frequencies, preconditioning procedures) and differences in model type and injury severity (e.g., NP depressurization). Time-dependent properties depend largely on NP status^69^, and pronounced degeneration may be necessary to detect measurable changes. Most viscoelasticity data originated from physical disruption or hybrid models, which may contribute to variability in the reported responses. The observed reductions in leakage and saturation pressure indicated impaired disc pressurization following injury, which may reduce the disc’s ability to maintain internal fluid pressurization^70^ ^71^.

Altogether, physical and biochemical disruption of IVD yielded non-uniform mechanical deterioration, stiffness, Young’s modulus, and RoM provide robust markers of degeneration, whereas viscoelasticity may remain largely unaffected. Species significantly moderated all five analyzed mechanical domains, but for most species, only a single study contributed data, making it impossible to separate species effects from study-specific factors. Nonetheless, pooling data across species is supported by prior research showing similar disc collagen content^12^ and comparable mechanical resistance when normalized by disc geometric parameters like height and area^16^ ^72^, though absolute lumbar RoMs are smaller in non-human species compared to human, particularly in flexion-extension^73^.

Further preclinical models should explicitly consider sex, IVDD induction method, and testing methodology effects, using standardized protocols, integrating mechanical and structural endpoints to improve data comparability and translational relevance. Despite statistical significance in pooled outcomes, wide prediction intervals crossing zero constrain generalizability.

### Morphological Changes following IVD injury

Consistent reductions in disc height and increases in degeneration grade were observed following IVD injury, in agreement with previous findings from in vivo animal models^6^ and clinical studies^74^. Disc height loss in rats was a reproducible marker of degeneration across studies. Degenerative grade exhibited a large pooled effect size, but the evidence was limited to the male rat subgroup and accompanied by high heterogeneity and evidence of publication bias. The substantial heterogeneity across studies may be due to differences in IVD injury models, spine levels, timepoints, and scoring systems. Standardized histological grading will be important to improve comparability cross models.

### Relationship Between Mechanical and Morphological Changes

Although data overlap was insufficient for direct statistical correlation, the observed pattern of concurrent mechanical impairment and structural degeneration supports their interdependency. Consistent changes in Young’s modulus, RoM, disc height, and degeneration grade reinforced the concept that morphological degeneration is accompanied by functional compromise, though the time course relationship could not be assessed due to the sparse distribution of timepoints.

## Limitations

The overall methodological quality of included studies was moderate, particularly limited by lacking sample size calculations and unclear or high risk of selection and performance bias due to poor methodological reporting. Diverse injury types, spine levels, injury severities, post-injury timepoints, mechanical testing directions, and outcome definitions contributed substantial heterogeneity. Pooling different testing directions under the same domain for meta-analysis may have obscured important directional differences, such as higher sensitivity of bending mechanics to NP pressurization compared to torsion^6^. Such methodology variation limits comparability across studies and generalizability of findings.

Use of adjacent IVDs or segments as controls could be problematic and may bias comparisons, because local IVD injury has been shown to result in degeneration in adjacent IVDs and structures^75–77^. Additionally, few studies reported both morphological and mechanical outcomes at the same IVD level, making it impossible to analyse the correlation between structural and functional degeneration.

Limited studies per moderator level reduced meta-regression analysis power and robustness. IVDD evaluation timepoints varied greatly across studies, ranging from one week to 18 months for mechanical outcomes. Rats were the only species included for morphological meta-analyses, assessed at one to 12 weeks post-injury. Pooling outcomes across disparate timepoints introduced heterogeneity and limited temporal interpretation. Moreover, different parameters may follow distinct time courses after IVD injury^50^ ^51^.

Evidence of publication bias was detected in several domains (e.g., stiffness, Young’s modulus, RoM, degeneration grade), suggesting the need for more inclusive reporting, including publication of negative results^78^. Future investigations should prioritize standardization of injury protocols and outcome measures, systematic integration of morphological and mechanical endpoints, and protocol pre-registration to enhance transparency and reproducibility.

## Conclusion

This systematic review and meta-analysis demonstrated that physical and/or biochemical disruption of the IVD in in vivo animal models leads to significant mechanical and morphological degeneration. Among the mechanical parameters examined, Young’s modulus appeared to be the most sensitive indicator of mechanical deficits, followed by stiffness and RoM, although no single parameter captures the full spectrum of functional decline following disc injury. These findings support both the biological relevance and mechanical validity of preclinical IVDD models and reinforce their value for advancing disc degeneration research.

## Supporting information

search strategy, supplementary figure 1

## Reference

1. Teraguchi M, Yoshimura N, Hashizume H, et al. Prevalence and distribution of intervertebral disc degeneration over the entire spine in a population-based cohort: the Wakayama Spine Study. Osteoarthritis Cartilage 2014;22(1):104–10. doi: 10.1016/j.joca.2013.10.019 [published Online First: 2013/11/19]

2. Diwan AD, Melrose J. Intervertebral disc degeneration and how it leads to low back pain. JOR Spine 2023;6(1):e1231. doi: 10.1002/jsp2.1231 [published Online First: 2023/03/31]

3. Colombini A, Lombardi G, Corsi MM, Banfi G. Pathophysiology of the human intervertebral disc. Int J Biochem Cell Biol 2008;40(5):837–42. doi: 10.1016/j.biocel.2007.12.011 [published Online First: 20071228]

4. Vergroesen P-PA, Kingma I, Emanuel KS, et al. Mechanics and biology in intervertebral disc degeneration: a vicious circle. Osteoarthritis and cartilage 2015;23(7):1057–70. doi: doi:10.1016/j.joca.2015.03.028

5. Desmoulin GT, Pradhan V, Milner TE. Mechanical Aspects of Intervertebral Disc Injury and Implications on Biomechanics. Spine 2020;45(8):E457–E64. doi: 10.1097/brs.0000000000003291

6. Iatridis JC, Michalek AJ, Purmessur D, Korecki CL. Localized Intervertebral Disc Injury Leads to Organ Level Changes in Structure, Cellularity, and Biosynthesis. Cell Mol Bioeng 2009;2(3):437–47. doi: 10.1007/s12195-009-0072-8 [published Online First: 2009/09/01]

7. Salzer E, Schmitz TC, Mouser VH, et al. Ex vivo intervertebral disc cultures: degeneration-induction methods and their implications for clinical translation. Eur Cell Mater 2023;45:88–112. doi: 10.22203/eCM.v045a07 [published Online First: 20230329]

8. Rivera Tapia ED, Meakin JR, Holsgrove TP. In-vitro models of disc degeneration - A review of methods and clinical relevance. J Biomech 2022;142:111260. doi: 10.1016/j.jbiomech.2022.111260 [published Online First: 20220817]

9. Poletto DL, Crowley JD, Tanglay O, et al. Preclinical in vivo animal models of intervertebral disc degeneration. Part 1: A systematic review. JOR Spine 2023;6(1):e1234. doi: 10.1002/jsp2.1234 [published Online First: 20221220]

10. O’Connell GD, Vresilovic EJ, Elliott DM. Comparison of animals used in disc research to human lumbar disc geometry. Spine (Phila Pa 1976) 2007;32(3):328–33. doi: 10.1097/01.brs.0000253961.40910.c1 [published Online First: 2007/02/03]

11. Elliott DM, Sarver JJ. Young investigator award winner: validation of the mouse and rat disc as mechanical models of the human lumbar disc. Spine (Phila Pa 1976) 2004;29(7):713–22. doi: 10.1097/01.brs.0000116982.19331.ea [published Online First: 2004/04/17]

12. Showalter BL, Beckstein JC, Martin JT, et al. Comparison of animal discs used in disc research to human lumbar disc: torsion mechanics and collagen content. Spine (Phila Pa 1976) 2012;37(15):E900–7. doi: 10.1097/BRS.0b013e31824d911c [published Online First: 2012/02/16]

13. Smit TH. The use of a quadruped as an in vivo model for the study of the spine - biomechanical considerations. Eur Spine J 2002;11(2):137–44. doi: 10.1007/s005860100346 [published Online First: 2002/04/17]

14. Jin L, Balian G, Li XDJ. Animal models for disc degeneration-an update. Histol Histopathol 2018;33(6):543–54. doi: 10.14670/Hh-11-910

15. Elmounedi N, Keskes H. Establishment of intervertebral disc degeneration models; A review of the currently used models. Journal of orthopaedics 2024;56:50–56. doi: 10.1016/j.jor.2024.05.020 [published Online First: 20240513]

16. Fusellier M, Clouet J, Gauthier O, et al. Degenerative lumbar disc disease: in vivo data support the rationale for the selection of appropriate animal models. Eur Cell Mater 2020;39:18–47. doi: 10.22203/eCM.v039a02 [published Online First: 2020/01/07]

17. Peng Y, Qing X, Shu H, et al. Proper animal experimental designs for preclinical research of biomaterials for intervertebral disc regeneration. Biomater Transl 2021;2(2):91–142. doi: 10.12336/biomatertransl.2021.02.003 [published Online First: 20210628]

18. Lee NN, Salzer E, Bach FC, et al. A comprehensive tool box for large animal studies of intervertebral disc degeneration. JOR spine 2021;4(2):e1162. doi: 10.1002/jsp2.1162

19. Perie DS, Maclean JJ, Owen JP, Iatridis JC. Correlating material properties with tissue composition in enzymatically digested bovine annulus fibrosus and nucleus pulposus tissue. Ann Biomed Eng 2006;34(5):769–77. doi: 10.1007/s10439-006-9091-y [published Online First: 2006/04/07]

20. Risbud MV, Shapiro IM. Role of cytokines in intervertebral disc degeneration: pain and disc content. Nat Rev Rheumatol 2014;10(1):44–56. doi: 10.1038/nrrheum.2013.160 [published Online First: 2013/10/30]

21. Martin JT, Gorth DJ, Beattie EE, et al. Needle puncture injury causes acute and long-term mechanical deficiency in a mouse model of intervertebral disc degeneration. Journal of orthopaedic research: official publication of the Orthopaedic Research Society 2013;31(8):1276–82. doi: 10.1002/jor.22355

22. Xiao F, Noort W, Lévénez J, et al. An in vivo rat lumbar spine instability model induced by intervertebral disc injury. bioRxiv: the preprint server for biology 2025:2025.05.14.653956. doi: 10.1101/2025.05.14.653956

23. Melrose J, Shu C, Young C, et al. Mechanical destabilization induced by controlled annular incision of the intervertebral disc dysregulates metalloproteinase expression and induces disc degeneration. Spine (Phila Pa 1976) 2012;37(1):18–25. doi: 10.1097/BRS.0b013e31820cd8d5 [published Online First: 2011/12/20]

24. Rousseau M-AA, Ulrich JA, Bass EC, et al. Stab incision for inducing intervertebral disc degeneration in the rat. Spine 2007;32(1):17–24. doi: 10.1097/01.brs.0000251013.07656.45

25. Mosley GE, Wang M, Nasser P, et al. Males and females exhibit distinct relationships between intervertebral disc degeneration and pain in a rat model. Sci Rep 2020;10(1):15120. doi: 10.1038/s41598-020-72081-9

26. Page MJ, McKenzie JE, Bossuyt PM, et al. The PRISMA 2020 statement: an updated guideline for reporting systematic reviews. Rev Esp Cardiol (Engl Ed) 2021;74(9):790–99. doi: 10.1016/j.rec.2021.07.010 [published Online First: 2021/08/28]

27. Bramer WM, Giustini D, de Jonge GB, et al. De-duplication of database search results for systematic reviews in EndNote. J Med Libr Assoc 2016;104(3):240–3. doi: 10.3163/1536-5050.104.3.014

28. Otten R, de Vries R, Schoonmade L. Amsterdam Efficient Deduplication (AED) method (Version 1): Zenodo; 2019 [Available from: https://zenodo.org/records/4544315.

29. Macleod MR, O’Collins T, Horky LL, et al. Systematic Review and Metaanalysis of the Efficacy of FK506 in Experimental Stroke. Journal of Cerebral Blood Flow & Metabolism 2005;25(6):713–21. doi: 10.1038/sj.jcbfm.9600064

30. Hooijmans CR, Rovers MM, de Vries RB, et al. SYRCLE’s risk of bias tool for animal studies. BMC Med Res Methodol 2014;14:43. doi: 10.1186/1471-2288-14-43 [published Online First: 20140326]

31. McGuinness LA, Higgins JPT. Risk-of-bias VISualization (robvis): An R package and Shiny web app for visualizing risk-of-bias assessments. Research Synthesis Methods 2020;n/a(n/a) doi: 10.1002/jrsm.1411

32. Guyatt GH, Oxman AD, Kunz R, et al. GRADE guidelines: 7. Rating the quality of evidence--inconsistency. J Clin Epidemiol 2011;64(12):1294–302. doi: 10.1016/j.jclinepi.2011.03.017 [published Online First: 20110731]

33. Zimmerman MC, Vuono-Hawkins M, Parsons JR, et al. The mechanical properties of the canine lumbar disc and motion segment. Spine (Phila Pa 1976) 1992;17(2):213–20. doi: 10.1097/00007632-199202000-00016 [published Online First: 1992/02/01]

34. Lü DS, Shono Y, Oda I, et al. Effects of chondroitinase ABC and chymopapain on spinal motion segment biomechanics. An in vivo biomechanical, radiologic, and histologic canine study. Spine (Phila Pa 1976) 1997;22(16):1828–34; discussion 34-5. doi: 10.1097/00007632-199708150-00006 [published Online First: 1997/08/15]

35. Wakano K, Kasman R, Chao EY, et al. Biomechanical analysis of canine intervertebral discs after chymopapain injection. A preliminary report. Spine 1983;8(1):59–68. doi: 10.1097/00007632-198301000-00011

36. Yuan Q, Du L, Xu H, et al. Autologous Mesenchymal Stromal Cells Combined with Gelatin Sponge for Repair Intervertebral Disc Defect after Discectomy: A Preclinical Study in a Goat Model. Frontiers in bioscience (Landmark edition) 2022;27(4):131. doi: 10.31083/j.fbl2704131

37. Gullbrand SE, Malhotra NR, Schaer TP, et al. A large animal model that recapitulates the spectrum of human intervertebral disc degeneration. Osteoarthritis Cartilage 2017;25(1):146–56. doi: 10.1016/j.joca.2016.08.006 [published Online First: 2016/08/30]

38. Detiger SE, Hoogendoorn RJ, van der Veen AJ, et al. Biomechanical and rheological characterization of mild intervertebral disc degeneration in a large animal model. J Orthop Res 2013;31(5):703–9. doi: 10.1002/jor.22296 [published Online First: 2012/12/21]

39. Wei Y, Tian Z, Tower RJ, et al. The Inner Annulus Fibrosus Encroaches on the Nucleus Pulposus in the Injured Mouse Tail Intervertebral Disc. American journal of physical medicine & rehabilitation 2021;100(5):450–57. doi: 10.1097/PHM.0000000000001575

40. Colloca CJ, Gunzburg R, Freeman BJ, et al. Biomechancial quantification of pathologic manipulable spinal lesions: an in vivo ovine model of spondylolysis and intervertebral disc degeneration. J Manip Physiol Ther 2012;35(5):354–66. doi: 10.1016/j.jmpt.2012.04.018

41. Fazzalari NL, Costi JJ, Hearn TC, et al. Mechanical and pathologic consequences of induced concentric anular tears in an ovine model. Spine (Phila Pa 1976) 2001;26(23):2575–81. doi: 10.1097/00007632-200112010-00010 [published Online First: 2001/11/29]

42. Latham JM, Pearcy MJ, Costi JJ, et al. Mechanical consequences of annular tears and subsequent intervertebral disc degeneration. *Clinical biomechanics (Bristol*, Avon) 1994;9(4):211–9. doi: 10.1016/0268-0033(94)90001-9

43. Lin H-J, Lin L-C, Hedman TP, et al. Exogenous Crosslinking Restores Intradiscal Pressure of Injured Porcine Intervertebral Discs: An In Vivo Examination Using Quantitative Discomanometry. Spine 2015;40(20):1572–7. doi: 10.1097/BRS.0000000000001089

44. Leckie AE, Akens MK, Woodhouse KA, et al. Evaluation of thiol-modified hyaluronan and elastin-like polypeptide composite augmentation in early-stage disc degeneration: comparing 2 minimally invasive techniques. Spine (Phila Pa 1976) 2012;37(20):E1296–303. doi: 10.1097/BRS.0b013e318266ecea [published Online First: 2012/07/10]

45. Buser Z, Kuelling F, Liu J, et al. Biological and biomechanical effects of fibrin injection into porcine intervertebral discs. Spine (Phila Pa 1976) 2011;36(18):E1201–9. doi: 10.1097/BRS.0b013e31820566b2 [published Online First: 2011/02/18]

46. Kaigle A, Ekstrom L, Holm S, et al. In vivo dynamic stiffness of the porcine lumbar spine exposed to cyclic loading: Influence of load and degeneration. Journal of Spinal Disorders 1998;11(1):65–70.

47. Kaigle AM, Holm SH, Hansson TH. 1997 Volvo Award winner in biomechanical studies. Kinematic behavior of the porcine lumbar spine: a chronic lesion model. Spine 1997;22(24):2796-806. doi: 10.1097/00007632-199712150-00002

48. Keller TS, Holm SH, Hansson TH, Spengler DM. 1990 Volvo Award in experimental studies. The dependence of intervertebral disc mechanical properties on physiologic conditions. Spine (Phila Pa 1976) 1990;15(8):751–61. doi: 10.1097/00007632-199008010-00004 [published Online First: 1990/08/01]

49. Crowley JD, Oliver RA, Wang T, et al. Lateral fenestration of lumbar intervertebral discs in rabbits: development and characterisation of an in vivo preclinical model with multi-modal endpoint analysis. European spine journal: official publication of the European Spine Society, the European Spinal Deformity Society, and the European Section of the Cervical Spine Research Society 2024(9301980) doi: 10.1007/s00586-024-08153-5

50. Fainor M, Orozco BS, Muir VG, et al. Mechanical crosstalk between the intervertebral disc, facet joints, and vertebral endplate following acute disc injury in a rabbit model. JOR spine 2023;6(4):e1287. doi: 10.1002/jsp2.1287

51. Calvo-Echenique A, Cegonino J, Correa-Martin L, et al. Intervertebral disc degeneration: an experimental and numerical study using a rabbit model. Medical & biological engineering & computing 2018;56(5):865–77. doi: 10.1007/s11517-017-1738-3

52. Leckie SK, Bechara BP, Hartman RA, et al. Injection of AAV2-BMP2 and AAV2-TIMP1 into the nucleus pulposus slows the course of intervertebral disc degeneration in an in vivo rabbit model. The spine journal: official journal of the North American Spine Society 2012;12(1):7–20. doi: 10.1016/j.spinee.2011.09.011

53. He S, Zhang Y, Zhou Z, et al. Similarity and difference between aging and puncture-induced intervertebral disc degeneration. Journal of orthopaedic research: official publication of the Orthopaedic Research Society 2022;40(11):2565–75. doi: 10.1002/jor.25281

54. Mosley GE, Hoy RC, Nasser P, et al. Sex Differences in Rat Intervertebral Disc Structure and Function Following Annular Puncture Injury. Spine 2019;44(18):1257–69. doi: 10.1097/BRS.0000000000003055

55. Boxberger JI, Auerbach JD, Sen S, Elliott DM. An in vivo model of reduced nucleus pulposus glycosaminoglycan content in the rat lumbar intervertebral disc. Spine (Phila Pa 1976) 2008;33(2):146–54. doi: 10.1097/BRS.0b013e31816054f8 [published Online First: 2008/01/17]

56. Norcross JP, Lester GE, Weinhold P, Dahners LE. An in vivo model of degenerative disc disease. Journal of orthopaedic research: official publication of the Orthopaedic Research Society 2003;21(1):183–8. doi: 10.1016/S0736-0266(02)00098-0

57. Xu H, Zhang Y, Zhang Y, et al. A novel rat model of annulus fibrosus injury for intervertebral disc degeneration. The spine journal: official journal of the North American Spine Society 2024;24(2):373–86. doi: 10.1016/j.spinee.2023.09.012

58. Thompson RE, Pearcy MJ, Barker TM. The mechanical effects of intervertebral disc lesions. Clin Biomech (Bristol, Avon) 2004;19(5):448–55. doi: 10.1016/j.clinbiomech.2004.01.012 [published Online First: 2004/06/09]

59. Elliott DM, Yerramalli CS, Beckstein JC, et al. The effect of relative needle diameter in puncture and sham injection animal models of degeneration. Spine 2008;33(6):588–96. doi: 10.1097/BRS.0b013e318166e0a2

60. Ulrich JA, Liebenberg EC, Thuillier DU, Lotz JC. ISSLS prize winner: repeated disc injury causes persistent inflammation. Spine (Phila Pa 1976) 2007;32(25):2812–9. doi: 10.1097/BRS.0b013e31815b9850 [published Online First: 2008/02/05]

61. Han B, Zhu K, Li F-C, et al. A simple disc degeneration model induced by percutaneous needle puncture in the rat tail. Spine 2008;33(18):1925–34. doi: doi:10.1097/BRS.0b013e31817c64a9

62. Zhang H, Yang S, Wang L, et al. Time course investigation of intervertebral disc degeneration produced by needle-stab injury of the rat caudal spine: laboratory investigation. J Neurosurg Spine 2011;15(4):404–13. doi: 10.3171/2011.5.Spine10811 [published Online First: 2011/07/05]

63. Michalek AJ, Iatridis JC. Height and torsional stiffness are most sensitive to annular injury in large animal intervertebral discs. Spine J 2012;12(5):425–32. doi: 10.1016/j.spinee.2012.04.001 [published Online First: 2012/05/26]

64. Meijer GJ, Homminga J, Veldhuizen AG, Verkerke GJ. Influence of interpersonal geometrical variation on spinal motion segment stiffness: implications for patient-specific modeling. Spine (Phila Pa 1976) 2011;36(14):E929–35. doi: 10.1097/BRS.0b013e3181fd7f7f

65. Liebsch C, Greiner-Perth AK, Vogt M, et al. Intervertebral disc degeneration, age, and sex affect the range of motion of the cervical spine. Sci Rep 2025;15(1):23540. doi: 10.1038/s41598-025-07182-4 [published Online First: 20250702]

66. Xiao F, Noort W, Lévénez J, et al. Mechanical instability of the lumbar spine following intervertebral disc injury: a comparison of two injury methods. Int Biomech 2025;12(1):67–80. doi: 10.1080/23335432.2025.2581469 [published Online First: 20251030]

67. Fujiwara A, Lim TH, An HS, et al. The effect of disc degeneration and facet joint osteoarthritis on the segmental flexibility of the lumbar spine. Spine (Phila Pa 1976) 2000;25(23):3036–44. doi: 10.1097/00007632-200012010-00011 [published Online First: 2001/01/06]

68. Tanaka N, An HS, Lim TH, et al. The relationship between disc degeneration and flexibility of the lumbar spine. Spine J 2001;1(1):47–56. doi: 10.1016/s1529-9430(01)00006-7 [published Online First: 2003/11/01]

69. Wang DL, Lai AL, Gansau J, et al. Ex vivo biomechanical evaluation of Acute lumbar endplate injury and comparison to annulus fibrosus injury in a rat model. J Mech Behav Biomed Mater 2022;131:9. doi: 10.1016/j.jmbbm.2022.105234

70. Appel Z, Michalek AJ. Enzymatic Digestion of the Intervertebral Disc Alters Intradiscal Injection and Leakage Mechanics. J Biomech Eng 2024;146(11) doi: 10.1115/1.4066071

71. Varden LJ, Nguyen DT, Michalek AJ. Slow depressurization following intradiscal injection leads to injectate leakage in a large animal model. JOR Spine 2019;2(3):e1061. doi: 10.1002/jsp2.1061 [published Online First: 20190907]

72. Beckstein JC, Sen S, Schaer TP, et al. Comparison of animal discs used in disc research to human lumbar disc: axial compression mechanics and glycosaminoglycan content. Spine (Phila Pa 1976) 2008;33(6):E166–73. doi: 10.1097/BRS.0b013e318166e001 [published Online First: 2008/03/18]

73. Alini M, Eisenstein SM, Ito K, et al. Are animal models useful for studying human disc disorders/degeneration? Eur Spine J 2008;17(1):2–19. doi: 10.1007/s00586-007-0414-y [published Online First: 2007/07/17]

74. Teichtahl AJ, Urquhart DM, Wang Y, et al. A Dose-response relationship between severity of disc degeneration and intervertebral disc height in the lumbosacral spine. Arthritis research & therapy 2015;17:297. doi: 10.1186/s13075-015-0820-1 [published Online First: 20151023]

75. Maas H, Noort W, Hodges PW, van Dieen J. Effects of intervertebral disc lesion and multifidus muscle resection on the structure of the lumbar intervertebral discs and paraspinal musculature of the rat. J Biomech 2018;70:228–34. doi: 10.1016/j.jbiomech.2018.01.004 [published Online First: 2018/02/06]

76. Kos N, Gradisnik L, Velnar T. A Brief Review of the Degenerative Intervertebral Disc Disease. Med Arch 2019;73(6):421–24. doi: 10.5455/medarh.2019.73.421-424

77. Constant C, Hom WW, Nehrbass D, et al. Comparison and optimization of sheep in vivo intervertebral disc injury model. JOR Spine 2022;5(2):e1198. doi: 10.1002/jsp2.1198 [published Online First: 20220422]

78. Joober R, Schmitz N, Annable L, Boksa P. Publication bias: what are the challenges and can they be overcome? J Psychiatry Neurosci 2012;37(3):149–52. doi: 10.1503/jpn.120065

