## Supplementary material for "Mechanical and morphological effects of intervertebral disc injury: a systematic review of in vivo animal studies": search strategy, supplementary figure 1

**Medline**

| **Search** | **Query** | **Results** |
| --- | --- | --- |
| #4 | #1 AND #2 AND #3 | **2,373** |
| #3 | exp "Biomechanical Phenomena"/ or exp "Mechanical Phenomena"/ or "Biophysics"/ or (("mech*" or "biomech*" or "biophysic*") adj3 ("phenom*" or "chang*" or "variation*" or "adaptation*" or "modif*" or "parameter*" or "propert*" or "charact*" or "respon*" or "alter*" or "effect*")).ti,ab,kf. or exp "Spine"/ or (("spine*" or "spinal*" or "vertebral column" or "Spinal column" or “intervertebral disk*” or “intervertebral disc*” or “inter vertebral disk*” or “inter vertebral disc*”) adj4 ("stiff*" or "hysteresis" or "stress-relaxation" or "range of motion" or "Viscoelasticity" or "neutral zone" or "creep")).ti,ab,kf. | 1,535,501 |
| #2 | (exp "Animals, Laboratory"/ or exp "Models, Animal"/ or ("rat" or "mouse" or "mice" or "rodent*" or "rabbit*" or "porcine" or "pig" or "bovine" or "sheep*" or "ovine" or "goat*" or "caprine" or "dog" or "beagle*" or "canine" or "baboon*" or "primate*" or "rhesus monkey*" or "non-human*" or "vertebrate*" or “cat” or “feline*”).ti,ab,kf.) not (exp "humans"/ not exp "Animals"/) | 4,146,401 |
| #1 | (exp "Intervertebral Disc"/ or (“back” or “lumba*” or “cervic*” or “spine*” or “spinal segment*”) adj3 ("injur*" or "damage*" or "stab*" or "lesion*" or "punct*" or "trauma*" or "wound*" or "damag*" or "disease" or pain or needle* or cut or cutting or scalpel or "knife" or "physical disruption").ti,ab,kf.) or (((“intervertebra*” or “inter-vertebra*” or “inter vertebra*”) adj1 (“disc*” or “disk*” or “segment*”).ti,ab,kf.)) adj3 ("injur*" or "damage*" or "stab*" or "lesion*" or "punct*" or "trauma*" or "wound*" or "damag*" or "disease" or "pain" or "needle*" or "cut" or "cutting" or "scalpel" or "knife" or "physical disruption").ti,ab,kf. | 152,814 |

**Embase**

| **Search** | **Query** | **Results** |
| --- | --- | --- |
| #5 | 4 not ("Conference Abstract" or "Conference Paper" or "Conference Review").pt. | **3,147** |
| #4 | 1 and 2 and 3 | 4,405 |
| #3 | exp "biomechanics"/ or exp "mechanics"/ or "Biophysics"/ or (("mech*" or "biomech*" or "biophysic*") adj3 ("phenom*" or "chang*" or "variation*" or "adaptation*" or "modif*" or "parameter*" or "propert*" or "charact*" or "respon*" or "alter*" or "effect*")).ti,ab,kf. or exp "Spine"/ or (("spine*" or "spinal*" or "vertebral column" or "Spinal column" or "intervertebral disk*" or "intervertebral disc*" or "inter vertebral disk*" or "inter vertebral disc*") adj4 ("stiff*" or "hysteresis" or "stress-relaxation" or "range of motion" or "Viscoelasticity" or "neutral zone" or "creep")).ti,ab,kf. | 4,287,609 |
| #2 | (exp "experimental animal"/ or exp "animal model"/ or ("rat" or "mouse" or "mice" or "rodent*" or "rabbit*" or "porcine" or "pig" or "bovine" or "sheep*" or "ovine" or "goat*" or "caprine" or "dog" or "beagle*" or "canine" or "baboon*" or "primate*" or "rhesus monkey*" or "non-human*" or "vertebrate*" or "cat" or "feline*").ti,ab,kf.) not (exp "human"/ not exp "Animal"/) | 5,100,841 |
| #1 | (((exp 'intervertebral disk'/ or ("back" or "lumba*" or "cervic*" or "spine*" or "spinal segment*").ti,ab,kf.) adj3 ("injur*" or "damage*" or "stab*" or "lesion*" or "punct*" or "trauma*" or "wound*" or "damag*" or "disease" or pain or needle* or cut or cutting or scalpel or "knife" or "physical disruption").ti,ab,kf.) or (("intervertebra*" or "inter-vertebra*" or "inter vertebra*") adj1 ("disc*" or "disk*" or "segment*")).ti,ab,kf.) adj3 ("injur*" or "damage*" or "stab*" or "lesion*" or "punct*" or "trauma*" or "wound*" or "damag*" or "disease" or "pain" or "needle*" or "cut" or "cutting" or "scalpel" or "knife" or "physical disruption").ti,ab,kf. | 208,736 |

**Web of Science**

| **Search** | **Query** | **Results** |
| --- | --- | --- |
| #4 | #1 AND #2 AND #3 | **1,229** |
| #3 | TS=((("mech*" or "biomech*" or "biophysic*") NEAR/3 ("phenom*" or "chang*" or "variation*" or "adaptation*" or "modif*" or "parameter*" or "propert*" or "charact*" or "respon*" or "alter*" or "effect*")) or (("spine*" or "spinal*" or "vertebral column" or "Spinal column" or “intervertebral disk*” or “intervertebral disc*” or “inter vertebral disk*” or “inter vertebral disc*”) NEAR/4 ("stiff*" or "hysteresis" or "stress-relaxation" or "range of motion" or "Viscoelasticity" or "neutral zone" or "creep"))) | 2,057,856 |
| #2 | (TS=((("Animal*” NEAR/3 (“Model*” or “Lab” or “Laborato*” or “Experiment*”)) or ("rat" or "mouse" or "mice" or "rodent*" or "rabbit*" or "porcine" or "pig" or "bovine" or "sheep*" or "ovine" or "goat*" or "caprine" or "dog" or "beagle*" or "canine" or "baboon*" or "primate*" or "rhesus monkey*" or "non-human*" or "vertebrate*" or “cat” or “feline*”))) | 5,277,161 |
| #1 | TS=("Intervertebral Disc" or ((“back” or “lumba*” or “cervic*” or “spine*” or “spinal segment*”) NEAR/3 ("injur*" or "damage*" or "stab*" or "lesion*" or "punct*" or "trauma*" or "wound*" or "damag*" or "disease" or “pain” or “needle*” or “cut” or “cutting” or “scalpel” or "knife" or "physical disruption")) or (((“intervertebra*” or “inter-vertebra*” or “inter vertebra*”) NEAR/1 (“disc*” or “disk*” or “segment*”))) NEAR/3 ("injur*" or "damage*" or "stab*" or "lesion*" or "punct*" or "trauma*" or "wound*" or "damag*" or "disease" or "pain" or "needle*" or "cut" or "cutting" or "scalpel" or "knife" or "physical disruption")) | 181,954 |

**Final Summary**

| Database | References Found | Final |
| --- | --- | --- |
| MEDLINE | 2373 |  |
| EMBASE | 3147 |  |
| WEB OF SCIENCE | 1229 |  |
| Total | 6,749 | 4,678 (2071 Duplicates) |

| **A1** | 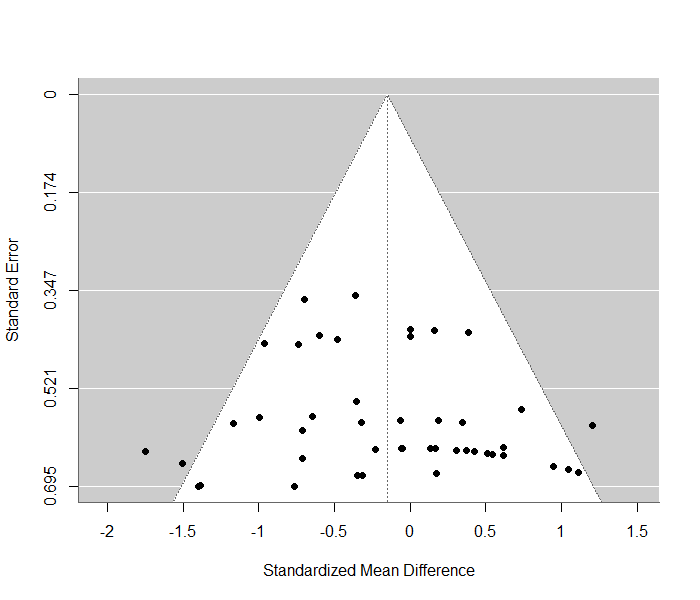 | **A2** | 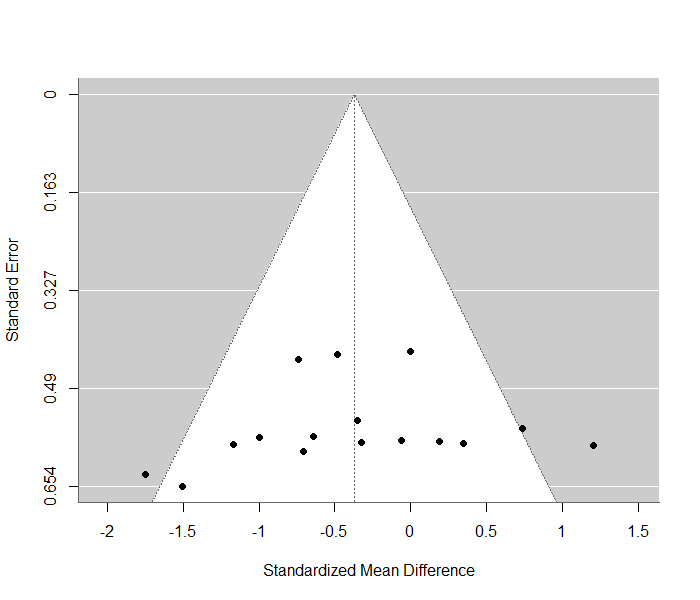 |
| --- | --- | --- | --- |
| **B** | 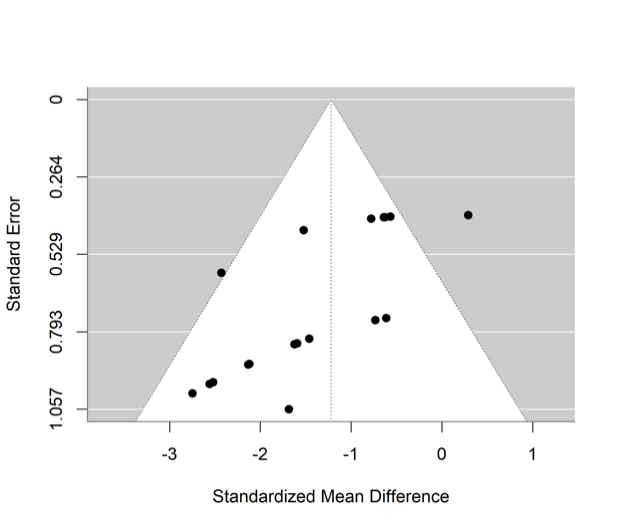 | **C** | 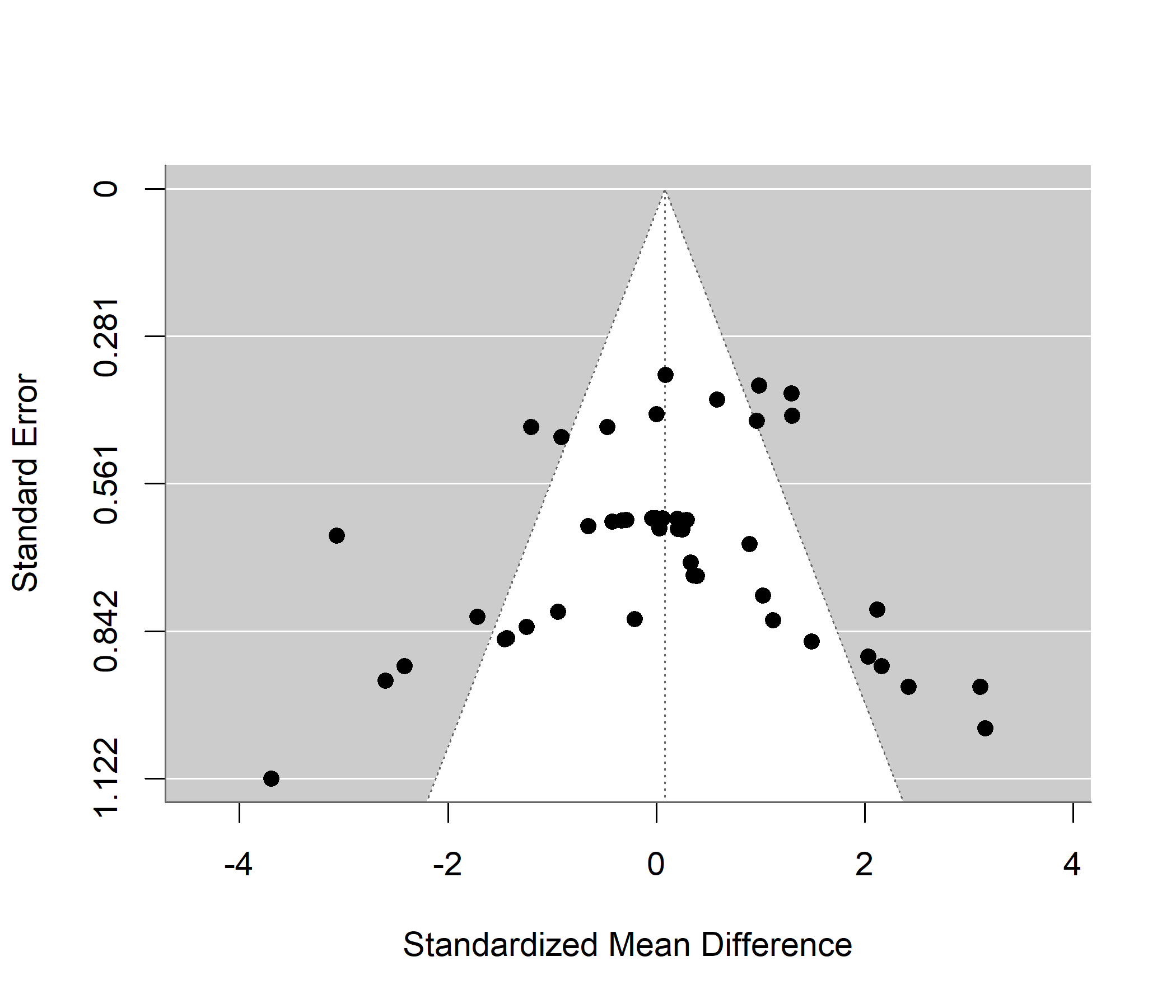 |
| **D** | 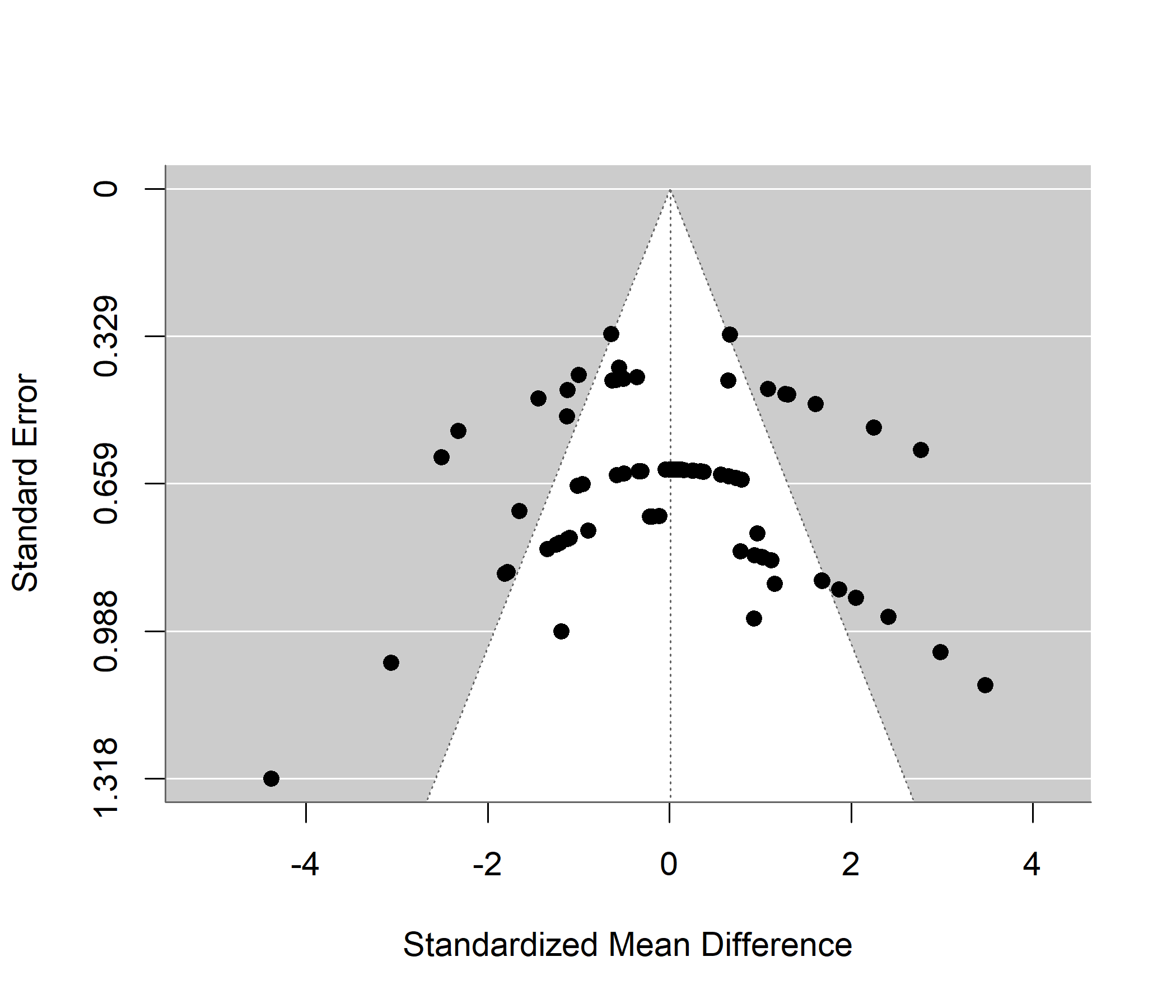 | **E** | 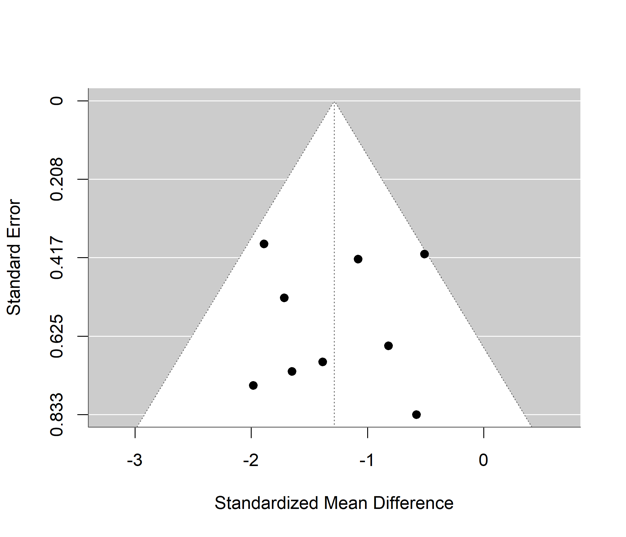 |
| **F** | 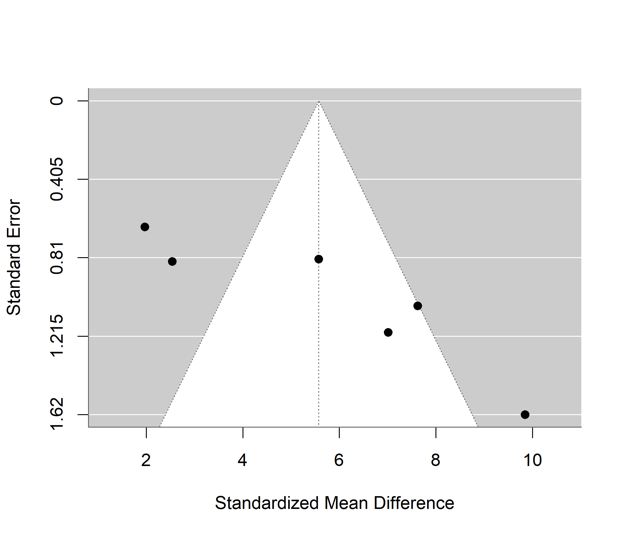 |  |  |
| **Figure S1**. Funnel plots of the Standardized Mean Difference (SMD) versus the standard error of the SMD for each of the included studies for (A1) stiffness-all models, (A2) stiffness-physical disruption models, (B) Young’s modulus, (C) range of motion, (D) viscoelasticity, (E) disc height, and (F) degeneration grade. | | | |
